# CTNND2 regulation by the SRGAP2 protein family links human evolution to synaptic neoteny

**DOI:** 10.1101/2022.09.13.507776

**Authors:** Nora Assendorp, Matteo Fossati, Baptiste Libé-Philippot, Eirini Christopoulou, Marine Depp, Roberta Rapone, Florent Dingli, Damarys Loew, Pierre Vanderhaeghen, Cécile Charrier

## Abstract

Human-specific genes are potential drivers of brain evolution. Among them, SRGAP2C has contributed to the emergence of features characterizing human cortical synapses, including their extended period of maturation. SRGAP2C inhibits its ancestral copy, the postsynaptic protein SRGAP2A; yet the synaptic molecular pathways differentially regulated in humans by SRGAP2 proteins remain largely unknown. Here, we identify CTNND2, a protein implicated in severe intellectual disability (ID) in the Cri-du-Chat syndrome, as an SRGAP2 effector. We demonstrate that CTNND2 slows down synaptic maturation and promotes neuronal homeostasis. During postnatal development, CTNND2 moderates neuronal excitation and excitability. In adults, it supports synapse maintenance. While CTNND2 deficiency is deleterious and results in the synaptic loss of SYNGAP1, another major ID-associated protein, the human-specific protein SRGAP2C enhances CTNND2 synaptic accumulation in human neurons. Our findings reveal that CTNND2 regulation by SRGAP2C contributes to synaptic neoteny in humans, and link human-specific and ID genes at the synapse.

## INTRODUCTION

During human evolution, cortical pyramidal neurons (CPNs) specialized to mature over longer time scales, establish a greater number of synaptic connections, and integrate more information than in other species^1–12^. These distinctive features are thought to contribute to higher cognitive and social abilities characterizing *Homo Sapiens*, such as symbolic thought, complex language, long-term planning and transfer of knowledge across generations^13^. Their alterations by genetic mutations affecting synaptic development are a major cause of neurodevelopmental disorders such as intellectual disability (ID), autism spectrum disorders (ASD) or schizophrenia^14,15^. Therefore, identifying the molecular intersections between human brain evolution and neurological disorders, and understanding their role in the fundamental mechanisms of synaptic development represents a major challenge.

Despite growing understanding of human divergence at the genomic level^16,17^, the impact of human-specific genetic innovations on synapse cell biology remains largely unexplored. So far, only one family of genes specifically duplicated in humans, *SRGAP2* (Slit-Robo Rho GTPAse-activating protein 2), has been shown to regulate synaptic development and cortical connectivity^18–21^. About 3.3-2.4 million years ago, at a time coinciding with the emergence of the genus *Homo*, two successive large segmental duplications generated human-specific copies of the ancestral gene *SRGAP2A,* which were named *SRGAP2B* and *SRGAP2C* ^22,23^*. SRGAP2A* (SRGAP2 in other mammalian species) encodes a postsynaptic protein highly conserved in mammals that promotes the maturation of excitatory and inhibitory synapses and limits their density^18,19^. *SRGAP2B* and *SRGAP2C* both encode truncated versions of SRGAP2A that interact with SRGAP2A and inhibit its function. However, SRGAP2B’s function is attenuated compared to SRGAP2C due to amino-acid substitutions^21^, and its expression is thought to be altered by deletions that appeared upstream of its gene since the duplication^22^. As a consequence, SRGAP2C is considered the primary human-specific copy of the SRGAP2 family. When expressed in mouse CPNs, SRGAP2C delays synaptic maturation and increases synaptic density, and thereby supports the emergence of features characterizing human CPNs^18,19^. So far, all identified functions of SRGAP2C are mediated by the inhibition of ancestral SRGAP2A^18,19,24^.

Here, we used *SRGAP2* duplications as an entry point to identify molecular pathways that shape the development of synapses and are uniquely regulated in humans due to SRGAP2A inhibition by SRGAP2B/C. Using a proteomic screen in maturing mouse synapses, we identify catenin delta-2 (CTNND2) as a major binding partner of SRGAP2 linked to synapse organization. CTNND2 is an ID/ASD-associated protein involved in cadherin-mediated trans-synaptic interactions^25^. Its hemizygous loss is associated with ID in the Cri-du-Chat syndrome^25,26^, which accounts for approximately 1% of patients with profound ID^27^. Point mutations in CTNND2 have also been implicated in severe forms of autism^28^ and epilepsy^29^. In mice, CTNND2 is involved in synaptic plasticity, memory and cortical function^30,31^ but its role in synaptic development remains poorly understood^28,31–33^. Using gain and loss of function experiments in mouse CPNs in vivo and xenotransplantation of cortical neurons derived from human pluripotent stem cells (hPSC), we demonstrate that CTNND2 moderates the pace of synaptic development in a way similar to what has been shown for SRGAP2C^19,24^. Furthermore, CTNND2 controls the ratio between excitation and inhibition as well as neuronal excitability in juvenile mice, and it is required for dendritic spine long-term maintenance in adults, highlighting its major role in maintaining neuronal integrity. At the molecular level, we show that CTNND2 is necessary for the synaptic accumulation of SYNGAP1, a major excitatory postsynaptic protein also linked to accelerated maturation in ID^34^. SRGAP2C enhances the synaptic enrichment of CTNND2, and CTNND2 overexpression is sufficient to compensate for deficits induced by SRGAP2C loss of function in human synapses. Our results provide an evolutionary conserved mechanism that determines the pace of synaptic maturation, and whose human-specific regulation contributes to human synaptic neoteny.

## RESULTS

### Identification of SRGAP2 network of interactions for synapse organization

How SRGAP2 proteins regulate synaptic development is poorly understood. Since SRGAP2C inhibits SRGAP2A^18,19,21^ (**Figure 1A**), the proteins associated with SRGAP2A are likely to be differentially regulated in human neurons. To map the interaction network of SRGAP2 (mouse ortholog of SRGAP2A) in maturing synapses, we performed a quantitative proteomic screen based on the co-immunoprecipitation of endogenous SRGAP2 and label-free liquid chromatography tandem mass spectrometry (LC–MS/MS) in synaptic fractions from postnatal day (P)15 mouse brains (i.e., during the period of synaptic maturation) (**Figure 1B-C, Table S1**). SRGAP2 interaction network was associated with human phenotypes including microcephaly, skull and brain size, neurodevelopmental delay and ASD, consistent with SRGAP2 implication in human brain evolution (**Figure 1D**). Moreover, synaptic gene ontology analysis using SYNGO^35^ highlighted a major implication of proteins involved in synapse organization, which primarily depends on postsynaptic scaffolds and trans-synaptic interactions^36^ (**Figure 1D-E**). Among all proteins in SRGAP2 network involved in synapse organization, we identified CTNND2 as the most prominent in terms of enrichment and estimated abundance (**Figure 1C, E**). CTNND2 is a brain-specific component of the cadherin-catenin adhesion complex involved in trans-synaptic interactions^37^. Its mutations are associated with ID in the Cri-du-Chat syndrome^25,26^ and severe forms of autism^28^. SRGAP2A and CTNND2 also co-immunoprecipitated in synaptosomes from the human neocortex (**Figure 1F**). Experiments in heterologous HEK cells revealed that CTNND2 could also form a complex with the human-specific protein SRGAP2C (**Figure 1G**). These results identify CTNND2 as a potential central target for human-specific regulations of synaptic development.

**Figure 1.**
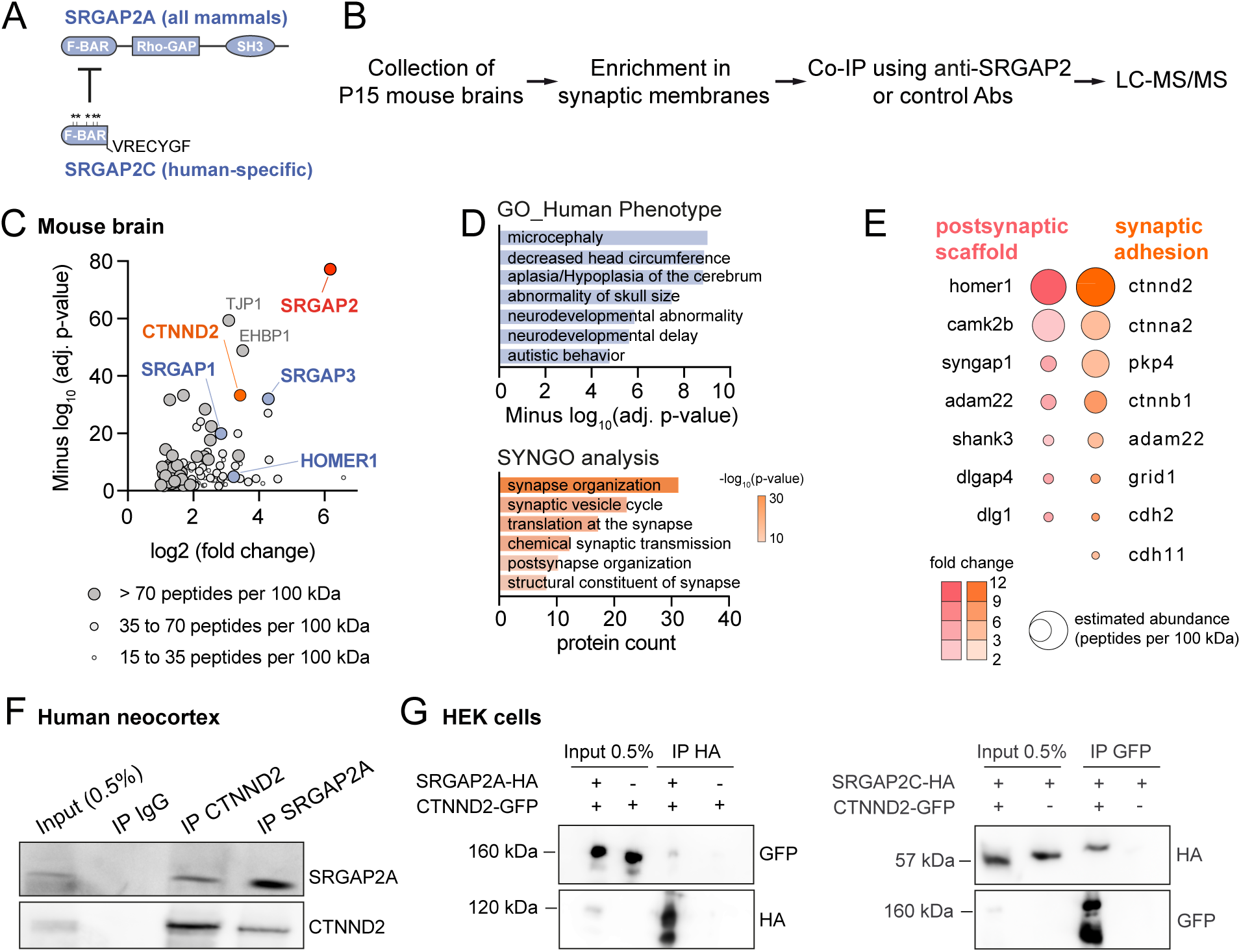
SRGAP2 protein interaction network at synapses. (A) Schematic of SRGAP2A and SRGAP2C protein domain composition and interaction. SRGAP2C lacks the last 49 amino-acids of the F-BAR domain of SRGAP2A. It contains a few point mutations (*) compared to SRGAP2A and ends with a unique stretch of 7 amino-acids. SRGAP2C inhibits SRGAP2A. (B) Experimental design. Co-IP, co-immunoprecipitation; LC-MS/MS, liquid chromatography followed by tandem mass spectrometry; Abs: antibodies. (C) Representation of SRGAP2 partners identified in LC-MS/MS depending on their enrichment, adjusted p-value, and estimated abundance. Only proteins with > 15 peptides per 100 kDa are represented. Data from 3 independent experiments. CTNND2 was one of the 10% most enriched proteins in SRGAP2 complexes, among known direct partners such as SRGAP3, SRGAP1 and HOMER1^18,61^. (D) Human phenotype and synaptic gene ontology (SYNGO) analysis of SRGAP2 interactome. (E) SRGAP2 partners involved in postsynaptic scaffolds and synaptic adhesion. (F) SRGAP2A and CTNND2 co-immunoprecipitate in synaptic fractions from human neocortex. (G) CTNND2-GFP co-immunoprecipitates with hemagglutinin (HA)-tagged SRGAP2A and SRGAP2C in HEK cells.

### CTNND2 limits spine density in juveniles but promotes spine maintenance in adults

Despite its implication in neurological disorders, the subcellular distribution and function of CTNND2 at synapses is poorly understood^37^. In mouse cortical neurons, we found that endogenous CTNND2 forms clusters associated with the vast majority (79 ± 4%) of excitatory synapses immunodetected with their core scaffolding protein PSD-95 (**Figure 2A**). To assess the role of CTNND2 at synapses, we performed sparse knockdown and knockout experiments in vivo in layer 2/3 (L2/3) CNPs of the mouse somatosensory cortex. Neural progenitors were transfected using in utero electroporation (IUE) at embryonic day (E) 15.5 with plasmids encoding either shRNAs or CAS9 and guide RNAs for CRISPR-mediated knockout, along with the fluorescent protein mVenus to visualize neuronal morphology (**Figure 2B; Figure S1-2**). We examined the consequences on dendritic spines as proxy for excitatory synapses and quantified their density in oblique apical dendrites using high-resolution confocal microscopy. In juvenile mice (P21), CTNND2-deficient neurons exhibited a higher density of dendritic spines than control neurons (knockdown: 115 ± 4% of shControl; **Figure 2C-D**; knockout: 116 ± 3% of Control; **Figure S2**). Normal spine density was rescued by co-electroporating shCtnnd2 with an shRNA-resistant *Ctnnd2* variant (96 ± 4% of control, see also **Figure S1**). On the other hand, CTNND2 overexpression strongly reduced spine density (59 ± 4% of control; **Figure 2C-D**). These effects were not associated with a significant change in dendritic spine morphology (not shown). These results demonstrate that CTNND2 regulates spine density in a dose-dependent manner and through cell-autonomous mechanisms. We next determined the time course of *Ctnnd2* knockdown consequences by examining two other developmental time points: P10, which corresponds to an early time point during spinogenesis, and P77, when mice have reached adulthood. In CTNND2-deficient neurons, spine density was not different from control at P10 but its rise between P10 and P21 was steeper than in control neurons (**Figure 2E-F**). In adult mice however, spine density in CTNND2-deficient neurons was lower than in control neurons (82 ± 4% of control, **Figure 2E-F),** consistent with previous work in a constitutive mouse model^31^. Thus, *Ctnnd2* deficiency has biphasic effect, leading first to an excess of dendritic spines followed by a precocious loss of synapses. To assess if spine loss in adult was a consequence of early defects during neuronal maturation, we used a conditional knockdown system. Expression of shCtnnd2 or shControl was induced after tamoxifen administration between P28 and P31 and spine density was quantified in P77 mice. Preserving CTNND2 expression during synaptic development did not prevent spine loss in adult mice (89 ± 3% of control, **Figure S3**), indicating independent functions of CTNND2 in dendritic spine development and long-term maintenance.

**Figure 2.**
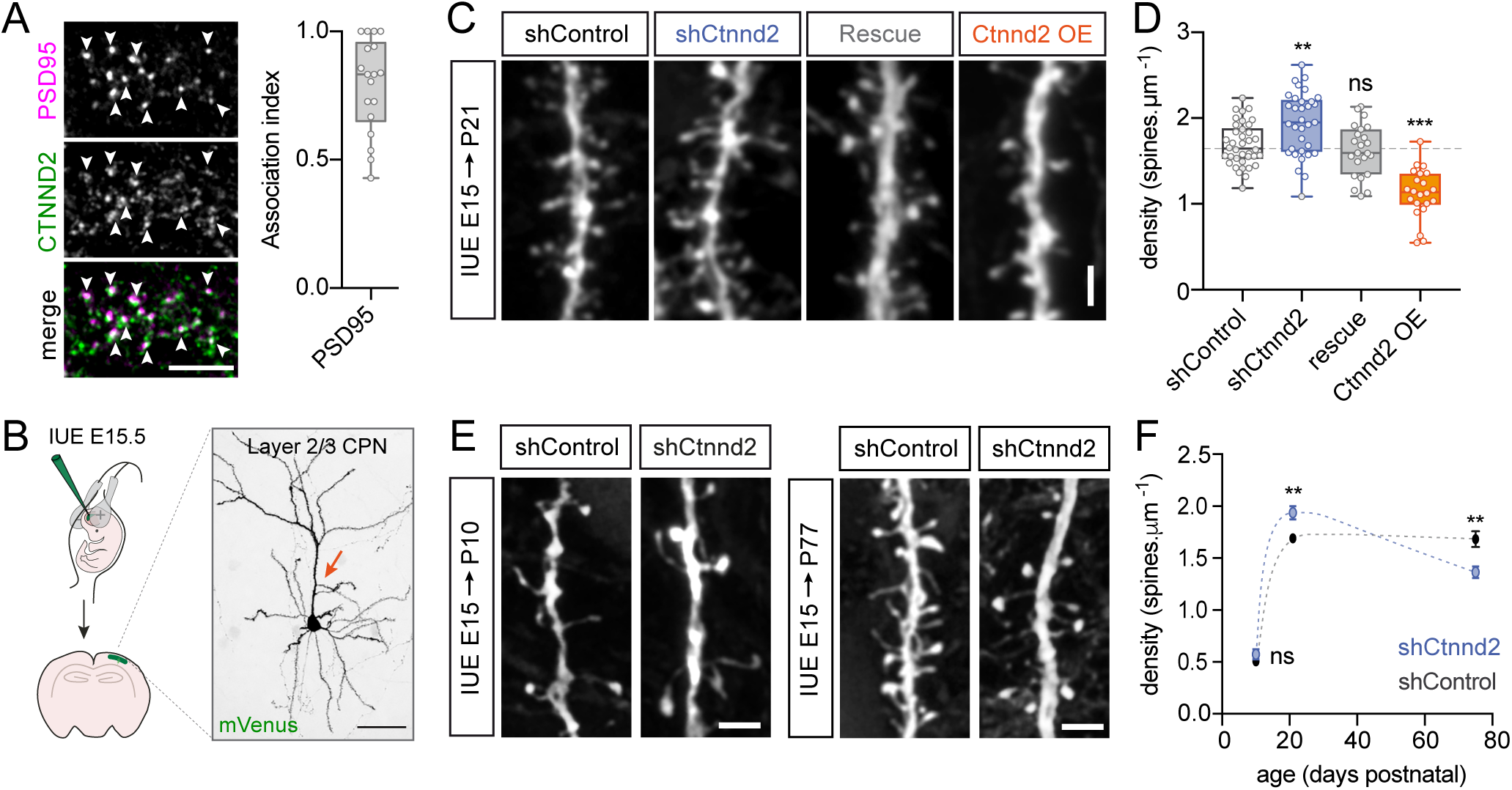
CTNND2 controls dendritic spine density and long-term maintenance. (A) Immunostaining of endogenous CTNND2 and PSD-95 in primary cortical neurons cultured for 18 days in vitro. Arrowheads points the association of CTNND2 and PSD-95 clusters. Scale bar: 5 μm. Quantification of association index: N = 18 cells. (B) Schematic illustration of in utero electroporation (IUE). IUE at embryonic day (E)15.5 allows the sparse targeting of L2/3 CPNs. Arrow in the example image indicates oblique apical dendrites. Scale bar: 50 μm. (C) Confocal images illustrating dendritic spines in neurons expressing shControl, shCtnnd2, shCtnnd2 together with an shCtnnd2-resistant CTNND2 mutant (CTNND2*, rescue), or overexpressing CTNND2 (OE) along with mVenus in juvenile mice (P21). Scale bar: 2 μm. (D) Quantifications of dendritic spine density in the conditions described above. N_shControl_ = 38 cells (6 mice); N_shCtnnd2_ = 33 (7); N_Rescue_ = 23 (5); N_Ctnnd2 OE_ = 22 (4). Dashed line indicates median of shControl condition. (E-F) Same as C-D in young (P10) and adult (P77) mice. P10: N_(control)_ = 21 (3), N_(shCtnnd2)_ = 31 (3); P21: N_(control)_ = 38 (6), N_(shCtnnd2)_ = 33 (7) (same as D); P77: N_(control)_ = 20 (2), N_(shCtnnd2)_ = 24 (4). Statistics: ns: p > 0.05; **: p < 0.01 and ***: p < 0.001, one-way ANOVA followed by Tukey’s multiple comparisons test (D) or Student t-test (F). Related to Figures S1-3.

### CTNND2 controls the timing of synaptic maturation

To assess the physiological consequences of CTNND2 deficiency during the period of synaptic development, we next performed whole-cell patch-clamp recording in acute brain slices. We compared excitatory synaptic transmission in electroporated CTNND2-deficient neurons with neighboring non-electroporated control neurons. In juvenile mice (P17-P19), *Ctnnd2* knockdown increased the frequency of miniature excitatory postsynaptic currents (mEPSCs) (140 ± 12% of control) without modifying their amplitude (108 ± 6% of control, **Figure 3A-B**, see also **Figure S4** for CTNND2 knockdown in postmitotic neurons). Conversely, CTNND2 overexpression strongly decreased mEPSC frequency (67 ± 4% of control, **Figure 3C-D**) These results are in line with our morphological analysis. Remarkably, CTNND2 deficiency also affected mEPSC frequency at earlier stages (P9-P12), before any detectable change in spine density (**Figure 3E-F**). Day-by-day analysis between P9 and P11 revealed that the developmental increase in mEPSC frequency occurred earlier and was more intense in CTNND2-deficient neurons than in control neurons, whereas it was hampered by CTNND2 overexpression (**Figure 3G-J**). The increased frequency of mEPSCs in young CTNND2-deficient neurons was associated with higher AMPA/NMDA ratio (P10-12: 164 ± 23% of control, **Figure 3K**), suggesting that CTNND2 is a synaptic activity-limiting protein that regulates the timing at which AMPA receptors are recruited at synapses for their functional maturation^38^. Together, these results demonstrate that the dosage of CTNND2 determines the pace of synaptic maturation.

**Figure 3.**
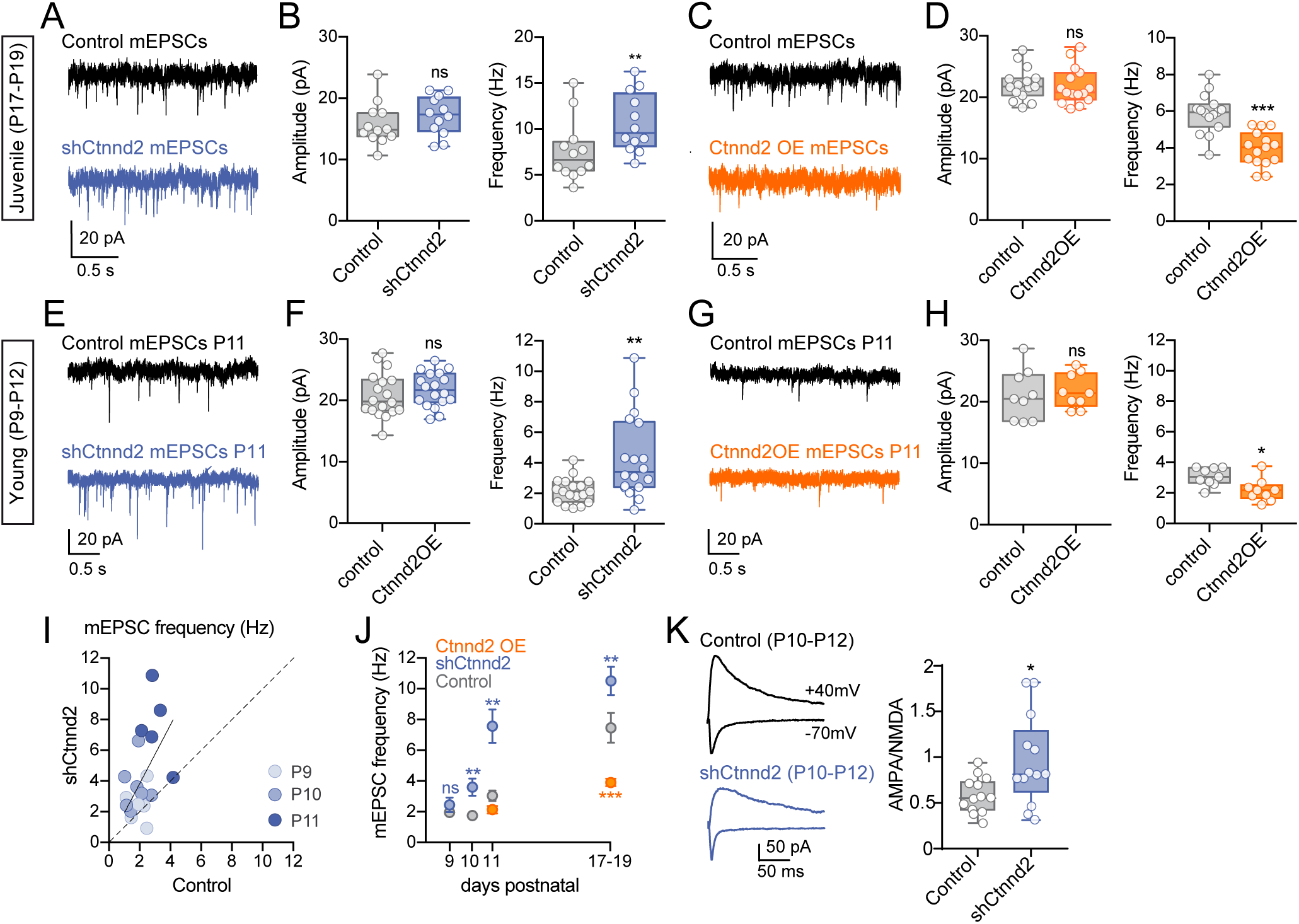
CTNND2 regulates the pace of synaptic maturation. (A-B) Representative traces and quantification of mEPSC recordings in acute brain slices from juvenile (P17-P19) mice. L2/3 CPNs expressing shCtnnd2 and tdTomato following IUE (blue) and neighboring non-electroporated control cells (gray) were compared. Data points in boxplots represent mean values for individual cells. N = 12 cells (10 mice) in each condition. (C-D) Same as A-B for control versus CTNND2-overexpressing neurons (OE). N = 15 (5) in each condition. (E-F) mEPSC analysis in control versus *Ctnnd2* knocked down neurons in young (P9-P11) mice. N = 18 (9) in each condition. (G-H) mEPSC analysis in control versus CTNND2-overexpressing neurons in young (P10-P12) mice. N = 9 (3) in each condition. (I) Scatterplots indicating mean mEPSC frequency for single pairs of neurons corresponding to shCtnnd2 and control neurons that were recorded from the same slice at indicated time points. (J) Mean mEPSC frequency at P9, P10, P11 and P17-19 (mean ± SEM) in the indicated conditions (same data as in A-H). (K) Representative traces and ratio of AMPA and NMDA receptor-mediated currents recorded at –70 mV and +40 mV, respectively. N = 13 (11) in each condition. Statistics: ns: p > 0.05; **: p < 0.01; *** p < 0.001, Student t-test (A-J), Mann-Whitney test (K). Related to Figure S4.

### CTNND2 limits excitation and excitability

We then tested how the increase in excitation in juvenile mice impacts the ratio between excitatory and inhibitory synaptic transmission (E/I ratio). To that aim, we placed a stimulation electrode approximately 100 μm above the cell body along the main apical dendrite (**Figure 4A**) and recorded evoked excitatory and inhibitory postsynaptic currents (eEPSC and eIPSC) by clamping the cell at –70 mV and +10 mV, respectively (**Figure 4B**). CTNND2 depletion strongly increased the E/I ratio compared to control neurons (185% of maximum current amplitude in control neurons and 152% of control synaptic charge over 50 ms, **Figure 4C**), uncovering a net excess of excitation in CTNND2-deficient L2/3 CPNs. In addition, CTNND2 deficiency increased intrinsic excitability. Although the resting membrane potential (Vrest) and spike threshold (Th) did not differ between control and CTNND2-deficient juvenile neurons (Vrest_Control_ = –78 ± 1 mV, Vrest_shCtnnd2_ = –76 ± 1 mV; Th_Control_ = –37 ± 1 mV, Th_shCtnnd2_ = –35 ± 1 mV), the membrane resistance of CTNND2-deficient neurons was higher (152 ± 11% of control), their input/output relationship stronger, their rheobase lower (73 ± 5% of control) and they fired action potential at a much higher frequency than control neurons (162 ± 8% of control for 500 pA current injection) (**Figure 4D-J**). By contrast, no difference was observed when non-electroporated control neurons were compared to neurons electroporated with a control shRNA (**Figure S5**). The increased excitability combined with the excess of excitation suggests the failure of autoregulatory and homeostatic mechanisms in juvenile CTNND2-deficient L2/3 CPNs.

**Figure 4.**
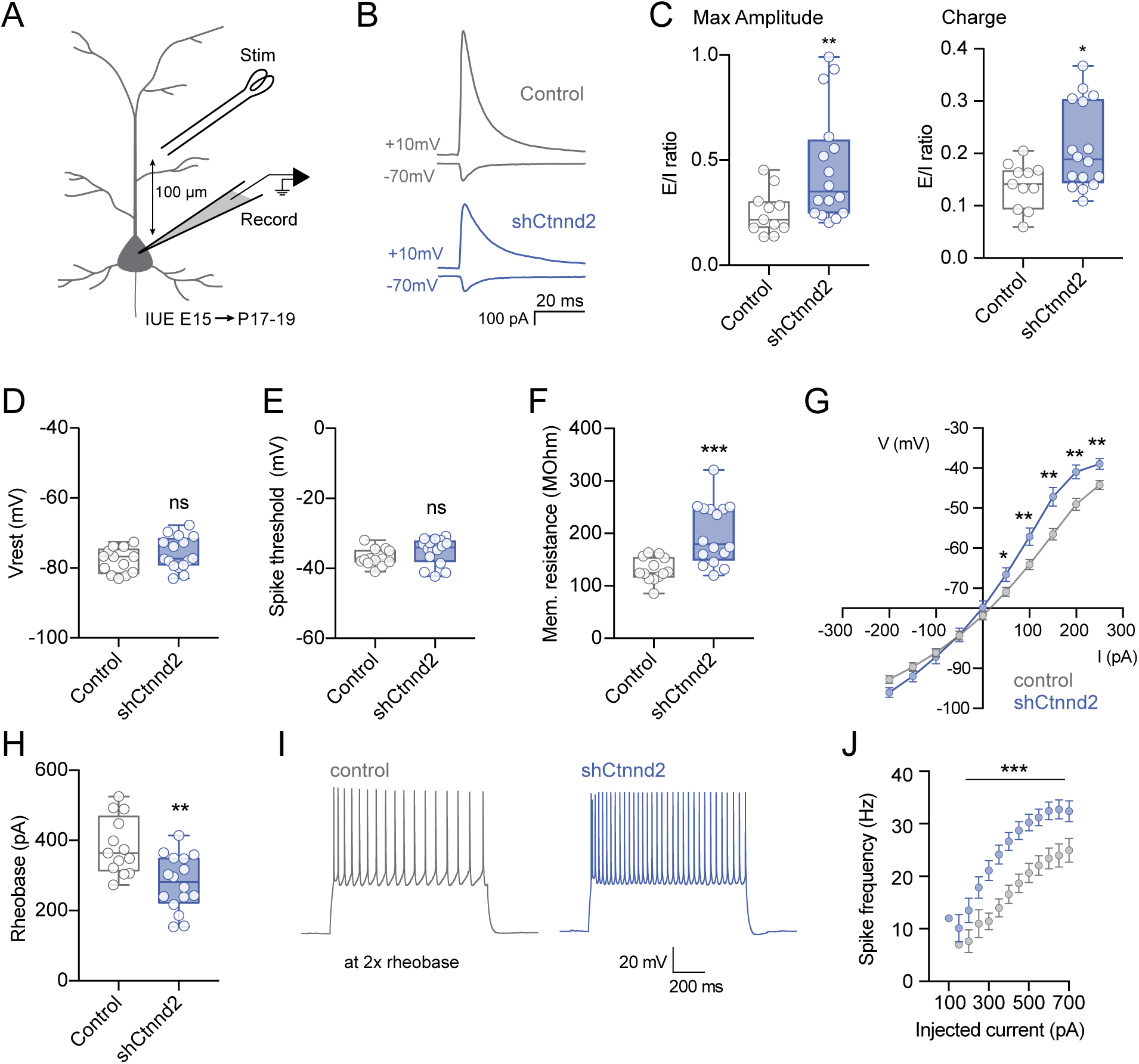
CTNND2 controls the E/I ratio and intrinsic excitability. (A) Evoked postsynaptic currents were recorded in L2/3 CPNs in acute slices from P17-P19 mice by placing a stimulation electrode approximately 100 μm above the cell body along the apical dendrite. (B) Representative traces of eIPSCs and eEPSCs in control and shCtnnd2-expressing neurons. (C) E/I ratio calculated using the maximum amplitude or the synaptic charge. N_control_ = 11 (6 mice) and N_shCtnnd2_ = 16 cells (10). (D-H) Quantification of intrinsic membrane properties in control and CTNND2-deficient L2/3 CPNs in acute slices from P17-19 mice. (I) Traces of action potentials evoked at twice the rheobase by 1s depolarizing current steps in whole-cell current-clamp recordings. (J) CTNND2-deficient neurons fire action potentials at higher frequency than control neurons. N_control_= 13 cells (5 mice), N_shCtnnd2_ = 16 (7). ns: p > 0.05; **: p < 0,01 and ***: p < 0,001, Student t-test. Related to Figure S5.

### CTNND2 is necessary for SYNGAP1 accumulation at excitatory synapses

To understand how CTNND2 regulates synaptic maturation, we performed a quantitative proteomic screen and identified CTNND2 protein interaction network at synapses in P15 mice, as described for SRGAP2 (**Figure 5A-B, Table S1**). SYNGO analysis of the most enriched partners highlighted their implication in synapse organization, trans-synaptic interactions, chemical synaptic transmission, regulation of neurotransmitter receptor level at synapses and regulation of plasma membrane potentials, consistent with our morphological and physiological results (**Figure 5C**). In addition to members of the cadherin/catenin superfamily (**Figure 5B,D**) and ion channels involved in the control of intrinsic excitability^8,39–41^ (**Figure 5D**), one of most prominent (>90 peptides per 100 kDa) partners of CTNND2 we identified was the excitatory postsynaptic protein SYNGAP1 (**Figure 5B,D**). SYNGAP1 is one of the most abundant proteins of excitatory postsynaptic densities (PSDs)^36^. Its haploinsufficiency accounts for up to 1% of non-syndromic ID in humans^34^. SYNGAP1 regulates excitatory synaptic transmission by competing with AMPA receptors for the binding to PSD95^34,42^. Its deficiency, like CTNND2 deficiency, accelerates excitatory synaptic maturation^43–45^. To assess a possible interplay between CTNND2 and SYNGAP1, we co-electroporated in utero SYNGAP1-GFP along with tdTomato and either shCtnnd2 or shControl, and we analyzed SYNGAP1 accumulation in individual dendritic spines along oblique apical dendrites of L2/3 CPNs at P21. CTNND2 deficiency led to a major decrease of SYNGAP1 level in spines (44 ± 9% of control) without change in the level of endogenous PSD-95 labelled with GFP-tagged FingRs^46^ (113 ± 24% of control) (**Figure 5E-F**, see also **Figure 7**). These results demonstrate that CTNND2 is necessary for the synaptic targeting of SYNGAP1 and the formation of an ID-associated protein complex that limits synaptic activity and delays synaptic maturation.

**Figure 5.**
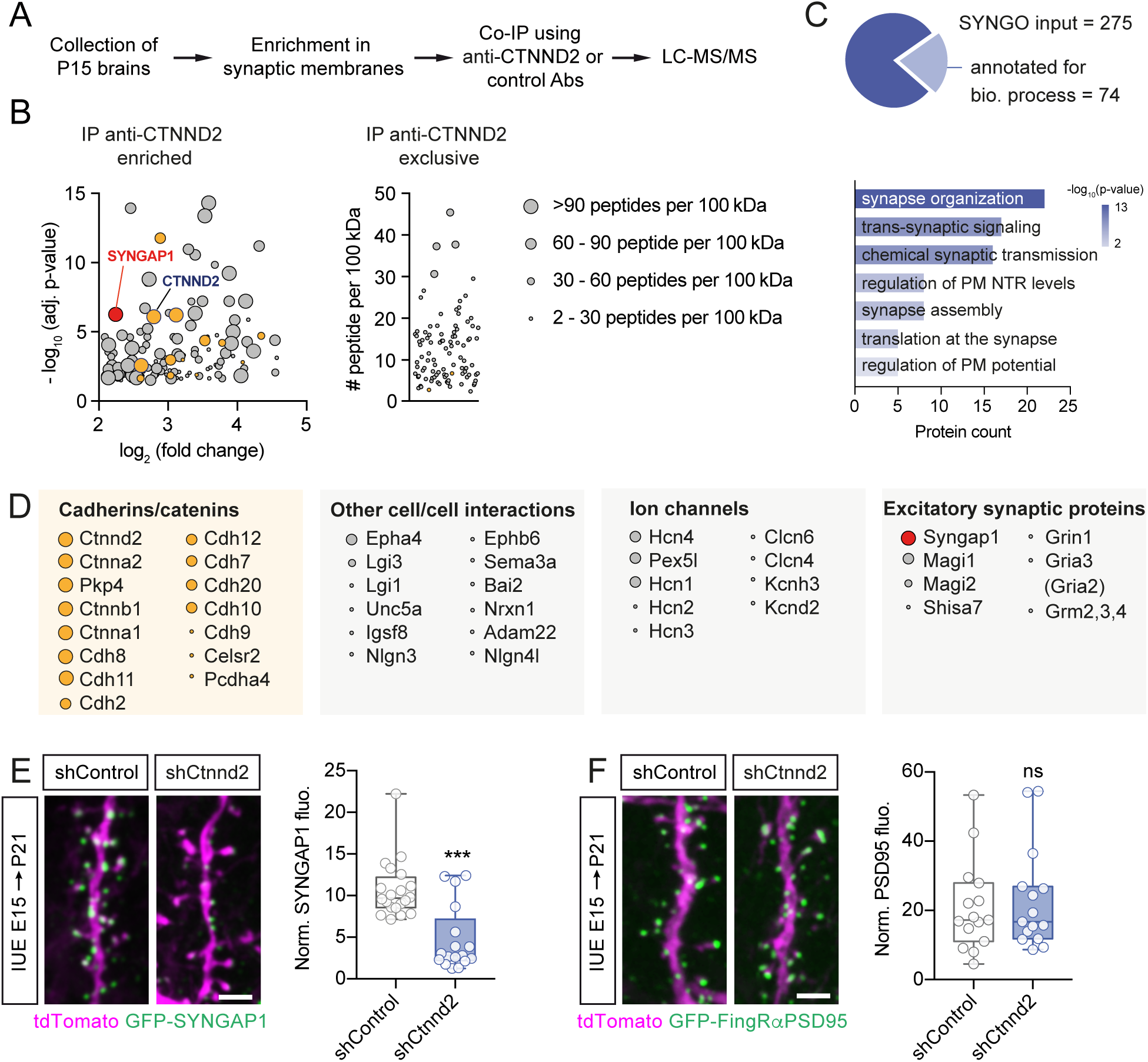
CTNND2 recruits SYNGAP1 to synapses. (A) Schematic of the experimental workflow. (B) Relative distribution and estimated abundance of proteins enriched in CTNND2 immunoprecipitates. Only proteins enriched by at least 4-fold, with at least 5 peptides, and whose p-value was < 0.05 based on three independent experiments are displayed. Cadherins and Catenins are shown in orange, SYNGAP1 is in red. (C) SYNGO analysis of CTNND2 partners. The pie chart indicates the proportion of CTNND2 identified partners annotated in SYNGO. (D) Classes of proteins and individual components standing out in CTNND2 protein complexes. Although not included in (B) due to stringent threshold, GluA2 was also detected in CTNND2 complexes (3-fold enrichment, p = 0.007, 32 peptides / 100 kDa). (E-F) Confocal images illustrating SYNGAP1 or PSD-95 puncta and quantification of their amount in dendritic spines. L2/3 CPNs were electroporated with tdTomato and SYNGAP1-GFP or GFP-tagged FingRs against PSD-95, together with shControl or shCtnnd2. E: N_shcontrol_ = 17, N_shCtnnd2_ = 20 cells (2 mice); F: N = 15 dendrites (4 mice) in each condition. Scale bar: 2 μm. Statistics: ns: p > 0.05; ***: p < 0,001 Mann-Whitney test.

### CTNND2 is necessary for human synaptic neoteny

We next sought to determine the role of CTNND2 in synaptic development in human neurons. We knocked down *CTNND2* and expressed EGFP in CPNs derived from hPSCs using lentivirus infection after 31 days in vitro, we selected for postmitotic neurons using DAPT and ARA-C and, after 44 days in vitro, we transplanted these neurons into the cerebral ventricles of P0 mice as previously described^43,47^ (**Figure 6A**). Human CPNs transplanted into the postnatal mouse brain have been shown to retain their species-specific pace of maturation, integrate into functional neural circuits and reach a higher morphological, physiological and transcriptomic level of maturation than neurons cultured in purely in vitro 2D or 3D assays^47–50^. We quantified the density of dendritic spines 2-, 6– and 10-months post-transplantation (MPT). *CTNND2* knockdown accelerated the formation of dendritic spines between 2 and 6 MPT, and spine density remained at a higher level than in control neurons 10 MPT (**Figure 6B-C**), indicating that the role of CTNND2 in regulating the pace of synaptic development is conserved across species. Remarkably, the effects of *CTNND2* knockdown in human neurons phenocopy those observed following *SRGAP2B/C* knockdown in similarly transplanted CPNs^24^. These results demonstrate that, like SRGAP2 duplications, CTNND2 contributes to synaptic neoteny in human neurons.

**Figure 6.**
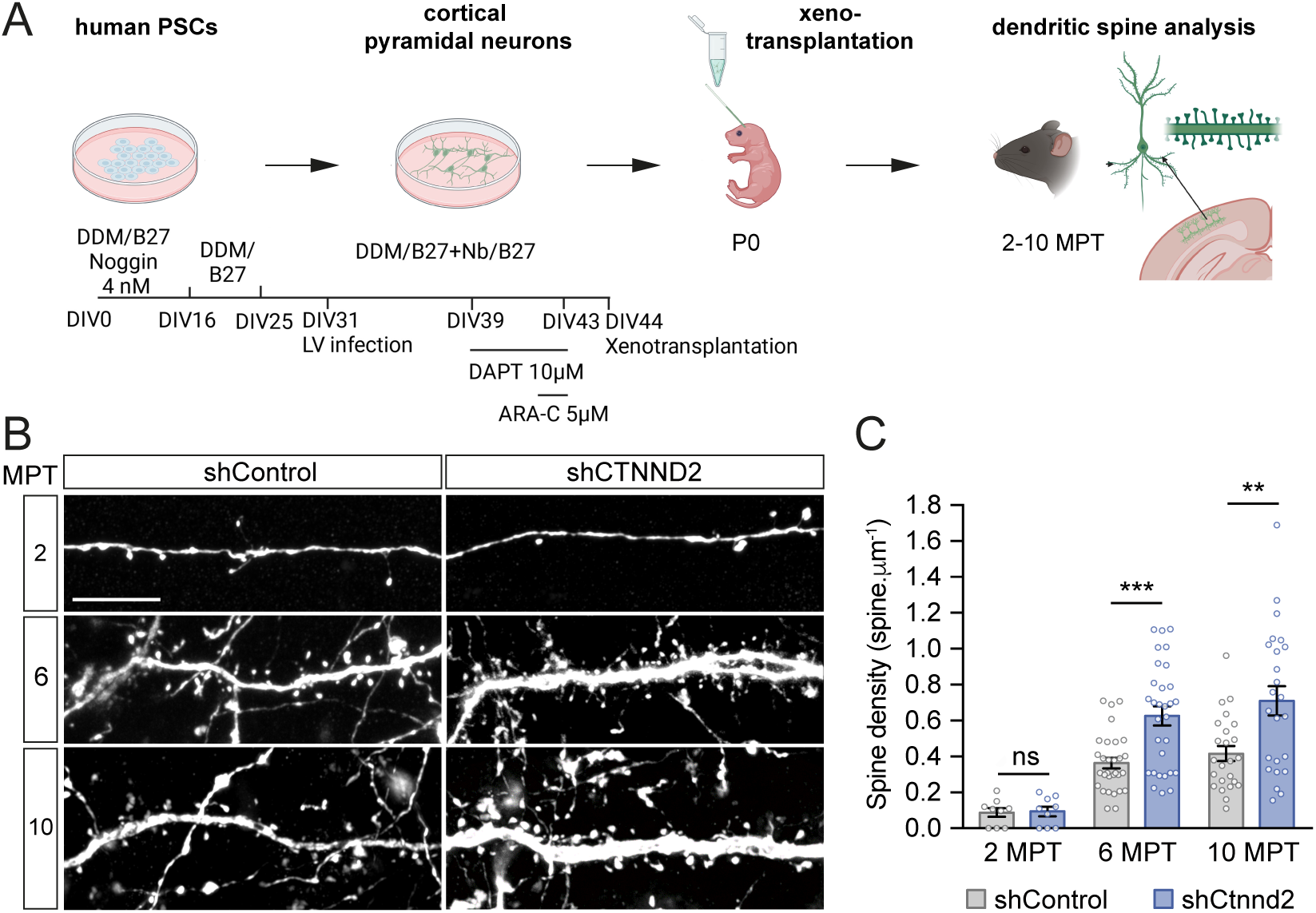
CTNND2 regulates dendritic spine development in human neurons. (A) Experimental design: human pluripotent stem cell (PSC)-derived neurons were infected in vitro with lentivirus (LV) expressing EGFP and control shRNAs or shRNAs targeting *CTNND2* transcripts, and then xenotransplanted in the mouse neonatal cerebral cortex. Morphological analysis was performed 2-10 months post-transplantation (MPT). Cartoons generated using Biorender.com. (B) Representative proximal dendritic branches of human PSC-derived neurons 2 to 10 MPT. Scale bar: 10 μm. (C) Quantification of dendritic spine density (mean ± SEM). N=9-30 neurons from 3-6 mice from 2 litters per stage per condition. Statistics: ns: p > 0.05; **: p < 0.01; ***: p < 0,001 Mann-Whitney test.

### CTNND2 is enhanced in human synapses by SRGAP2C

We next addressed whether and how SRGAP2 proteins regulate CTNND2. Because the pace of neuronal and synaptic development in human neurons is a major limitation to molecular investigations of neuron cell biology, we first expressed SRGAP2C in mouse L2/3 CPNs (**Figure 7**). In agreement with our previous work^18^, juvenile mouse L2/3 CPNs expressing SRGAP2C following IUE exhibited lower synaptic levels of PSD-95 labelled with GFP-FingRs (75 ± 5% of control), reflecting a delay in postsynaptic scaffold assembly and synaptic maturation. By contrast, SRGAP2C expression strongly enhanced the enrichment of CTNND2-GFP in individual dendritic spines (146 ± 8% of control, **Figure 7A-B**). We then isolated synaptosomes from the cortex of P15 wild-type and *Srgap2* heterozygous mice^19^, a model recapitulating the human situation in which SRGAP2A is partially inactivated by SRGAP2B/C. CTNND2 level in P15 cortical synaptosomes was about three times higher in *Srgap2* heterozygous mice than in wild-type mice (**Figure 7C-D**). Altogether, these data indicate that SRGAP2A inhibition by SRGAP2C is sufficient to increase CTNND2 level at synapses.

**Figure 7.**
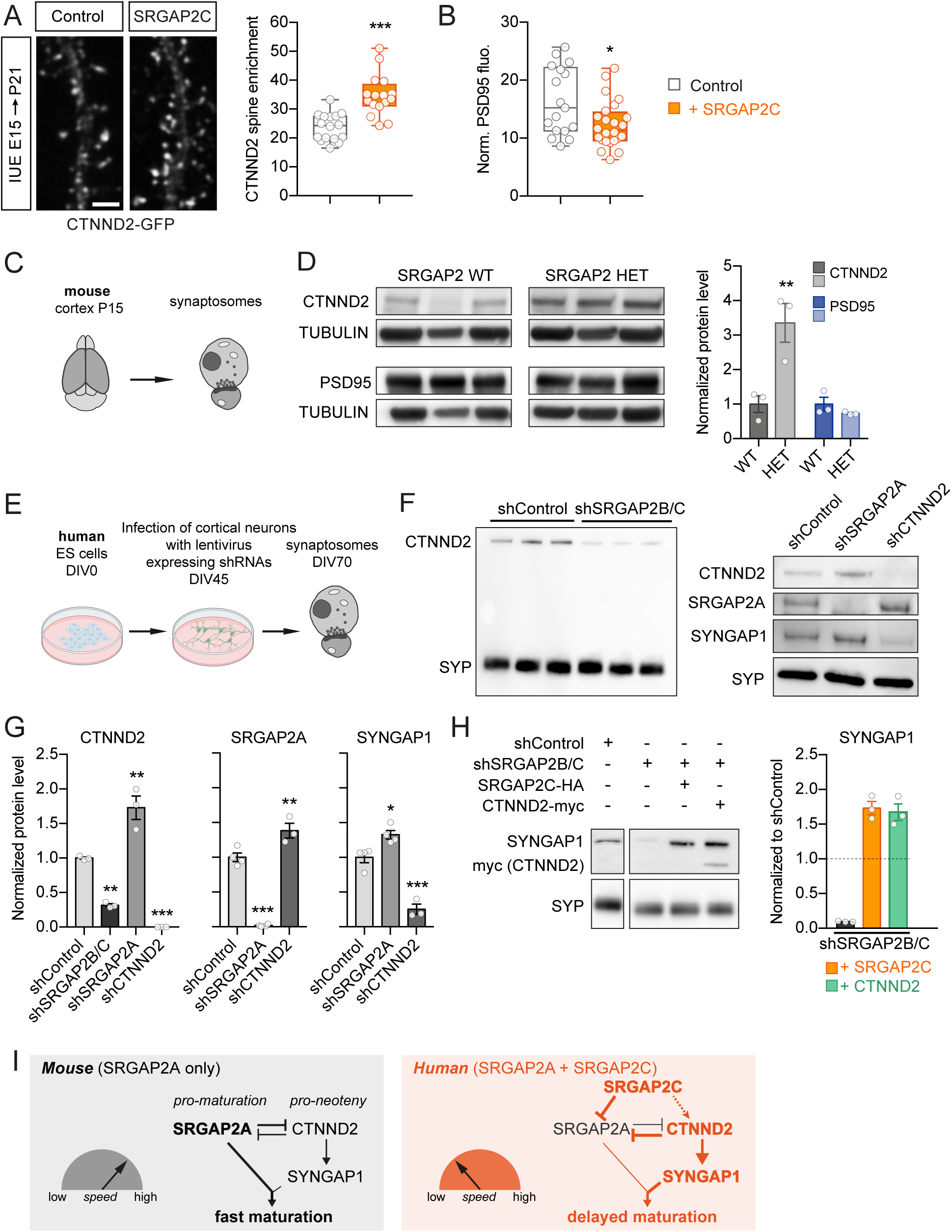
The SRGAP2 protein family regulates CTNND2 synaptic accumulation. (A-B) Confocal images and quantification of CTNND2 and PSD-95 amounts in individual spines in control and SRGAP2C-expressing mouse P21 neurons following mouse IUE at E15.5. CTNND2 was tagged with GFP and endogenous PSD-95 was labelled with GFP-tagged FingRs. CTNND2: N_control_ = 16 cells (3 mice), N_SRGAP2C_ = 16 (3); PSD-95: N_control_ = 17 (6), N_SRGAP2C_ = 24 cells (7). Scale bar: 2 μm. Statistics: ns: *: p < 0.05; ***: p < 0,001 Mann-Whitney test. (C-D) Normalized amount of CTNND2 and PSD-95 in synaptosomes prepared from cortices of P15 wild-type (WT) and SRGAP2 heterozygous (HET) mice measured in western blot (N = 3). (E) Experimental design: human cortical neurons were differentiated from embryonic stem (ES) cells and transduced with lentiviruses expressing shRNAs after 45 days in vitro (DIV). Proteins were extracted from synaptosomes at DIV70 and assessed in western blot. Cartoons generated using Biorender.com. (F-G) Representative western blot and quantification of SRGAP2A, CTNND2, synaptophysin (SYP) and SYNGAP1 in synaptosomal protein extracts from human neurons in the indicated conditions (N = 3-4). ANOVA followed by Tukey’s multiple comparison test. (H) SYNGAP1 level in synaptosomal extracts from human neurons infected with lentiviruses to knockdown SRGAP2C and overexpress SRGAP2C-HA or CTNND2-myc. Data are expressed as fraction of the level of SYNGAP1 in shControl condition. The shControl lane on the right is from the same blot. N=3. (I) Model of the regulation of synaptic maturation by CTNND2 and SRGAP2 proteins in mouse and human neurons. In mice, SRGAP2A promotes synaptic maturation by limiting the synaptic accumulation of CTNND2, and consequently of SYNGAP1. In humans, SRGAP2C partially inhibits SRGAP2A. This leads to an increase in CTNND2 synaptic enrichment, which further lowers SRGAP2A synaptic level and recruits more SYNGAP1 to synapses, providing a powerful mechanism to delay synaptic maturation in human neurons. Related to Figure S6.

To interrogate more directly the regulation between SRGAP2 proteins and CTNND2 in human neurons, we differentiated cortical neurons from hPSCs in vitro. Neurons were selected using L1CAM antibodies to remove progenitors and non-neuronal cells from the cultures, infected after 45 days in vitro with lentivirus expressing shRNAs and cultured for 70 days (see methods), at which point synaptosomes were prepared for western blot analysis (**Figure 7E**). We observed bidirectional regulation of CTNND2 by SRGAP2 proteins: CTNND2 level in synaptosomes strongly decreased after *SRGAP2B/C* knockdown (32 ± 4% of shControl, **Figure 7F-G**) and increased following *SRGAP2A* inactivation (173 ± 17% of shControl). SRGAP2A level in human synaptosomes was also increased by *CTNND2* knockdown (138 ± 11% of shControl, **Figure 7F-G**), suggesting cross-inhibition between CTNND2 and SRGAP2A.

Last, we tackled the regulation of SYNGAP1 by CTNND2 and SRGAP2 proteins in human neurons. In agreement with our results in mice, CTNND2 was necessary for the accumulation of SYNGAP1 in human synaptosomes (**Figure 7F**). SYNGAP1 level in human synaptosomes increased following *SRGAP2A* inactivation, albeit to a lesser extent than CTNND2 (**Figure 7F).** Consistently, it also decreased after *SRGAP2B/C* knockdown (**Figure 7G**, see also^24^). Since (1) CTNND2 is necessary for SYNGAP1 accumulation at synapses; and (2) SRGAP2A associates with CTNND2 but not SYNGAP1 (**Figure S6**), we tested the hypothesis that CTNND2 mediates the regulation of SYNGAP1 by SRGAP2C in human neurons. In synaptosomes from human *SRGAP2B/C* knocked down neurons, the expression of a shRNA-resistant SRGAP2C cDNA and CTNND2 overexpression both rescued SYNGAP1 level in a similar way (**Figure 7H**). Collectively, these results demonstrate that CTNND2 is specifically regulated in human neurons by SRGAP2 duplications, which in turn impacts the synaptic recruitment of SYNGAP1 and the pace of synaptic maturation in human cortical neurons (**Figure 7I**).

## DISCUSSION

Understanding the cellular and molecular basis of the distinctive traits of the human brain represents a major challenge in neuroscience with broad implications for medicine. Because the development of the human brain is protracted and spans over two decades, the mechanisms underlying the specialization of human cortical synapses have remained largely intractable. Here, we mapped the synaptic protein interaction network of SRGAP2, one of the few proteins specifically duplicated in humans and the only one implicated in synaptic development so far^18–20^. Our results reveal that CTNND2, one of SRGAP2’s most prominent partners, sets the pace of synaptic maturation and that its regulation by the human-specific protein SRGAP2C underlies synaptic neoteny in human neurons. Our results provide molecular insights into the regulation of CTNND2 by SRGAP2C. They also highlight that CTNND2 is necessary for the synaptic accumulation of SYNGAP1 and the formation of a complex of ID/ASD-associated proteins uniquely regulated in humans that shapes the developmental trajectory of synapses. Changes in the timing of developmental events (“heterochrony”), including onset, duration or speed, are thought to be a major driver of interspecies phenotypic variation across evolution^51^ and are associated with neurodevelopmental disorders in humans^15,52,53^. Our results uncover a molecular intersection between evolutionary innovations, human-specific regulations and disorder-associated dysfunctions in neocortical synapses.

So far, the role of CTNND2 at synapses has remained debated. Previous studies suggested that CTNND2 loss of function either decreases^28^ or increases^33^ the density of dendritic spines in young neurons, or that CTNND2 is not involved in synaptic development but rather in spine maintenance in adult mice^31^. The reasons for these inconsistencies are unclear. However, several studies used a “N-term” mouse model^28,30,31,33^, often referred to as “KO”, which may not fully recapitulate the consequences of CTNND2 loss of function. Indeed, this mouse model still expresses a truncated protein corresponding to the first ≈ 445 amino-acids of CTNND2 that contains most of the residues mutated in severe autism^28,30^. To clarify the role of CTNND2 at synapses, we performed knockdown, rescue, overexpression and CRISPR/CAS9-mediated knockout experiments with complementary approaches (high-resolution microscopy, slice electrophysiology, biochemistry and proteomics) and models (mice, hPSC-dervied neurons, xenotransplantation), and analysis at different developmental time points (young, juvenile and adults). Our results reveal that the dosage of CTNND2 at synapses controls the timing of synaptic maturation, with CTNND2 loss of function leading to earlier, faster and more intense increase in mEPSC frequency and dendritic spine density during postnatal development, and CTNND2 overexpression having opposite effects. These changes begin at a time coinciding with the opening of the critical period of plasticity in L2/3 CPNs^54^ and may have severe consequences in the context CTNND2 mutations in ID in the Cri-du-Chat syndrome and ASD.

In addition to the timing of synaptic maturation, we show that CTNND2 controls the level of excitation and intrinsic excitability in juvenile mice as well as the long-term maintenance of dendritic spines in adults. This suggests that the human-specific regulation of CTNND2 may provide an evolutionary advantage beyond its contribution to synaptic neoteny. In adults, the precocious loss of synapses in CTNND2-deficient neurons is consistent with the progressive retraction of dendrites and dendritic spines as well as with deficits in cortical responsiveness and cognition in mice carrying a CTNND2 partial truncation^31^. These alterations could reflect early neurodegeneration, a hypothesis supported by evidence that presenilin-1 mutations implicated in familial Alzheimer’s disease increase the proteolytic processing of CTNND2^31,55–57^. Conversely, mice expressing higher levels of CTNND2 exhibit better object recognition, better sociability, and lower anxiety than wild type mice^58^. Future experiments will determine whether CTNND2 regulation by SRGAP2C confers some degree of protection against synapse loss during aging.

Our results indicate that a balance between CTNND2 and SRGAP2A controls the timing of synaptic maturation. This balance is skewed by SRGAP2C in human neurons toward more CTNND2 (this manuscript) and less SRGAP2A^19,24^, which delays synaptic maturation and supports neoteny. The higher spine over shaft enrichment of CTNND2 in SRGAP2C-expressing neurons suggests a redistribution of CTNND2 in neurons. At synapses, SRGAP2A and CTNND2 may compete for interaction with common synaptic partners, including SRCIN1, CTNNA2, CEP170B, PKP4, cadherins, ADAM22, ARHGAP32 or AKAP10 (Table S1). However, since SRGAP2A is a Rho-GTPAse activating protein for Rac1^59^, CTNND2 synaptic level may also depend on RAC1 signaling and the contribution of other mechanisms, such as transcription, protein synthesis or protein stability cannot be excluded. Furthermore, while SRGAP2A inhibition is sufficient to enhance the synaptic level of CTNND2, CTNND2 interacts with SRGAP2C in heterologous cells, raising the possibility that SRGAP2C also directly regulates CTNND2 (**Figure 7I**).

Our results also shed a new light on the biology of postsynaptic scaffolds in the pathophysiology of ID/ASD. We found that CTNND2 is necessary for the synaptic accumulation of SYNGAP1, which is also regulated by SRGAP2 proteins (see also^24^). Accordingly, CTNND2 and SYNGAP1 are both ID/ASD-associated proteins whose inactivation have similar outcomes characterized by biphasic perturbations with early hyperexcitation and faster synaptogenesis, followed by adult cortical hypofunction in the mouse (this study and^15^). Moreover, a sticking acceleration of dendritic spine formation was found in xenotransplanted human CPNs following CTNND2 (this study) and SYNGAP1 disruption^24,43^. Therefore, SYNGAP1 deficit likely contributes to the pathogenesis of ID in the Cri-du-Chat syndrome, and CTNND2 upregulation could mitigate ID in patients with SYNGAP1 haploinsufficiency, opening new avenues for drug development in these disorders. Remarkably, CTNND2 accumulation in excitatory PSDs is regulated by SHANK3^60^, a prototypical and conserved postsynaptic scaffolding protein associated with Phelan-McDermid syndrome and ASD^15^. Together, these data raise fundamental questions on the hinge between the assembly of the postsynaptic scaffold and the recruitment of activity-limiting proteins to fine-tune the timing of synaptic maturation across species.

In the human neocortex, synaptic development is extremely protracted but the metronome remains poorly understood. We have previously shown that the human-specific protein SRGAP2C delays synaptic development in part by regulating its parental copy SRGAP2A and decelerating the assembly of the HOMER1-based excitatory postsynaptic scaffold^18^. Our current results uncover cross-inhibition between the pro-maturation protein SRGAP2A and the pro-neoteny factor CTNND2. The inhibition of SRGAP2A and the enhancement of CTNND2 synaptic enrichment by SRGAP2C, which in turn recruits more SYNGAP1 to synapses, provides a powerful mechanism for prolonging the period of synaptic maturation in human neurons. Collectively, our results provide a new understanding of how synaptic development is adjusted in humans at the core of postsynaptic machineries to generate species-specific physiological and functional properties. They highlight the link between human-specific and AD/ASD genes at the synapse and open new perspective to decipher the contribution of synaptic regulations specific to humans to the pathogenesis of neurological disorders.

## ACKNOWLEDGMENTS

We thank all the members of the Charrier lab and Mariano Casado for discussion. We thank the acute transgenesis facility, the imaging facility and the animal facility of IBENS for excellent support. We thank D. Arnold, B. Rico, C. Cepko, F. Polleux, F. Zhang, X. Morin, W.L. Jin and J. Godin for sharing reagents. This work was supported by INSERM, CNRS, ENS, the Agence Nationale de la Recherche (ANR-13-PDOC-0003 and ANR-17-ERC3-0009 to CC), the European Research Council (ERC starting grant SYNPATH to CC and ERC Adv Grant GENEVOCORTEX to PV), Labex Memolife, Fondation pour la Recherche Médicale (to NA and DL) and Région Ile-de-France (to DL), the EOS Programme, ERANET NEURON, the Belgian FWO, the Generet Foundation, and the Belgian Queen Elizabeth Foundation (to PV). BLP was supported by a postdoctoral Fellowship of the FWO (12V1219N).

## AUTHOR CONTRIBUTIONS

Conceptualization: NA, CC; Methodology: NA, MF, CC, FD, DL, BLP, PV; Investigation and formal analysis: NA, MF, MD, CC, RR, FD, BLP, PV; Visualization: NA, CC; Funding acquisition: CC, PV; Project administration: CC; Supervision: CC, PV; Writing original draft: NA, CC.

## STAR METHODS

### RESOURCE AVAILABILITY

#### Lead contact

Further information and requests for resources and reagents should be directed to the lead contact Cécile Charrier.

#### Materials availability

Data and reagents are available upon request.

#### Data and code availability

All data reported in this paper will be shared by the lead contact upon request. This paper does not report original code.

The proteomic datasets generated have been deposited on PRIDE (accession numbers are listed in the key resources table).

Any additional information required to reanalyze the data reported in this paper is available from the lead contact upon request.

## EXPERIMENTAL MODEL AND SUBJECT DETAILS

### Animals

All mice were handled according to EU regulations (directive 2010/63, decree 2013-118). For all experiments except xenotransplantations, authorization references were #1530-2015082611508691v3 and #29476 2021020311595454v5, delivered by the French Ministry of Research after evaluation by the Comité d’Ethique en Experimentation Animale n°005. For xenotransplantation in mice, experiments were performed with the approval of the KU Leuven Committee for animal welfare (protocol 2018/030). Mouse housing, breeding and experimental handling were performed according to the ethical guidelines of the Belgian Ministry of Agriculture in agreement with European community Laboratory Animal Care and Use Regulations (86/609/CEE, Journal officiel de l’Union européenne, L358, 18 December 1986).

All animals were housed under standard conditions (12 h light:12 h dark cycles) with food and water ad libitum. The day of birth was defined as P0. In utero electroporations were performed on pregnant Swiss females at E14.5-15.5 (Janvier labs) and electroporated offspring was used for experiments at varying postnatal timepoints from P9 to P77. Primary cultures were prepared from timed pregnant C57BL/6J or Swiss mice at E17.5-E18.5 (Janvier labs). The SRGAP2 mouse strain (B6;129P2-Srgap2Gt(XH102)Byg/Mmcd) has been described before^19^. *Rag2*^-/-^ mice used for xenotranplantations were purchased from Jackson Labs and bred and maintained in local facilities. Xenotransplantations were performed at P0 and analysis were performed 2 and 10 months later. A total of 26 mice of both sexes were used for xenotransplantation analysis. In all experiments, data were collected without consideration for the sex of the animals.

### Human tissue collection and preparation

The study on research involving human subjects was approved by the Ethics Committee Research of University Hospitals Leuven (UZ Leuven) (reference S61186). Prior to surgery, written informed consent was obtained. Information related to sex, age, race, socioeconomic status, cause of the surgery, and medication were retrieved from the medical report of the patients. Human cortical samples were resected from the cortex during neurosurgery. Samples were collected at the time of surgery, immerged in ice-cold ACSF (NaCl 126 mM, NaHCO_3_ 26 mM, D-glucose 10 mM, MgSO_4_ 6 mM, KCL 3 mM, CaCl_2_ 1 mM, NaH_2_PO4 1 mM, 295-305 mM, pH adjusted to 7.4, with 5% CO_2_/95% O_2_), transferred immediately to the laboratory, flash frozen in an interval of 5-10 minutes and stored at –80°C. Samples used in this study correspond to the healthy part of the resection from a 34-year-old female (temporal lobe epilepsy), 22-year-old female (frontal lobe epilepsy), 45-year-old female (temporal lobe epilepsy). These biopsies were also used in^24^.

### Cell lines and neuronal differentiation

HEK293T (CRL-1573 from ATCC; RRID:CVCL_0045) cells were cultured according to suggested protocols. Cells were maintained in DMEM (GIBCO) supplemented with 10% fetal bovine serum (GIBCO) and 1% Penicillin-Streptomycin (GIBCO) at 37°C, 5% CO2. They were passaged by trypsin/EDTA digestion (GIBCO) upon reaching confluency with a maximum of 25 passages. Human ESC H9 have been described previously^62^. Cells were maintained on irradiated mouse embryonic fibroblasts (MEF) in the ES medium until the start of cortical differentiation. Cortical differentiation from human ESC was performed as described previously^50^ with some modifications^47,63^. On DIV−2, ESCs were dissociated using Stem-Pro Accutase (Thermo Fisher Scientific, Cat#A1110501) and plated on matrigel-(hES qualified matrigel BD, Cat#354277) coated dishes at low confluency (5,000–10,000 cells/cm^2^) in MEF-conditioned hES medium supplemented with 10 μM ROCK inhibitor (Y-27632; Merck, Cat#688000). On DIV0 of the differentiation, the medium was changed to DDM^64^, supplemented with B27 devoid of Vitamin A (Thermo Fisher Scientific, Cat#12587010) and 100 ng/ml Noggin (R&D systems, Cat#1967-NG), and the medium was changed every 2 days until DIV6. From DIV6, the medium was changed every day until DIV16. After DIV16, the medium was changed to DDM, supplemented with B27 (DDM/B27), and changed every day. At DIV25, the progenitors were dissociated using Accutase and cryopreserved in mFreSR (StemCell Technologies, Cat#05855). Differentiated cortical cells were validated for neuronal and cortical markers by immunostaining using antibodies for TUBB3 (BioLegend, Cat#MMS-435P), TBR1 (Abcam, Cat#ab183032), CTIP2 (Abcam, Cat#ab18465), FOXG1 (Takara, Cat#M227), SOX2 (Santa Cruz, Cat#sc-17320), FOXP2 (Abcam, Cat#ab16046), SATB2 (Abcam, Cat#ab34735).

## METHOD DETAILS

### DNA constructs

#### Plasmids for protein expression

All plasmids for protein expression *in utero* have a pCAG backbone driving protein expression under the CAG promotor. pCAG SRGAP2A-HA and pCAG SRGAP2C-HA were previously described ^18,19^. For pCAG CTNND2-EGFP, human CTNND2 cDNA was obtained from Horizon (Clone IMAGE 40080647; Insert sequence: BC111837) subcloned by PCR and inserted in pCAG between HindIII and AgeI; pCAG CTNND2-Myc was created through digestion of pCAG CTNND2-EGFP between Age1 and Notl to remove the GFP and insertion of the DNA cassette containing 3xMYC with compatible ends; For rescue experiments, four silent point mutations were introduced in *ctnnd2* (c3242a_a3243t_g3244c_c3245t) to resist to shRNA-mediated knockdown (mutant named *ctnnd2*\*). For pCAG SYNGAP-EGFP, SYNGAP1 was amplified from cDNA (Horizon; ORFeome Collab. Hs SYNGAP1 ORF w/o Stop Codon; Clone IMAGE: 100015293; insert sequence: BC148357) and inserted into pCAG between BsrGI and NotI. pCAG_PSD95.FingR-eGFP-CCR5TC was a gift from Don Arnold (Addgene plasmid #46296; http://n2t.net/addgene:46296; RRID:Addgene_46296;^46^. All constructs were verified by DNA sequencing.

#### Plasmids for shRNA and CRISPR/CAS9

To express shRNAs, we used the previously described pH1SCV2 and pH1SCTdT2 vectors^18,19^ in which the H1 promoter drives the expression of the shRNA and a CAG promoter that of either mVenus (pH1SCV2) or TdTomato (pH1SCTdT2). mVenus was used for dendritic spine analysis and pH1SCTdT2 was used in patch clamp experiments. In some experiments (Figure S4), the shRNA was introduced in the vector pNeuroD2-miR-30a-shRNA (gift from Juliette Godin) between XhoI and EcoRI to restrict the expression of the shRNA to postmitotic neurons. The control shRNA (shControl) was described previously^19^. The shRNAs against *ctnnd2/CTNND2* were designed to inactivate both the mouse and human genes. The following seed sequences were inserted into the shRNA vectors: shCtnnd2#1 5’-GCAGTGAGATCGATAGCAAGA –3’ and shCtnnd2#2 5’-GGGAAATGATCAGCCTCAAAG –3’. Both shRNAs were used in spine density analysis at P21 and no difference was observed between the them. The mutant *ctnnd2\** is resistant against shCtnnd2#2. If not otherwise indicated shCtnnd2#2, which targets both mouse and human *CTNND2* was used in experiments. For conditional knock-down experiments, shControl or shCtnnd2 were inserted into pCAG-mir30 (Addgene plasmid #14758; ^65^) between EcoRI and XhoI. Then, the miR-30a-shRNA cassette was then amplified and inserted into pDIO-DSE-mCherry-PSE-MCS (gift from Beatriz Rico, Addgene plasmid #129669; http://n2t.net/addgene:129669; RRID:Addgene_129669;^66^) between PmeI and EcoRV. For Crispr-Cas9 knockout experiments, gRNAs were designed using the prediction software https://portals.broadinstitute.org/gpp/public/analysis-tools/sgrna-design and inserted in pX330-U6-Chimeric_BB-Cbh-hSpCas9 (gift from Feng Zhang, Addgene plasmid #42230; http://n2t.net/addgene:42230; RRID:Addgene_42230;^67^). The sequence of the gRNA targeting *ctnnd2* was GGAGCGATGCAAGCTTGGCT (in exon 3). For knockdown with lentivirus, we used a lentiviral vector (pLV) expressing shRNAs under a H1 promoter and GFP under a synapsin promoter, as previously described^68^ (see also^24^). The seed sequences were 5’-GCCACTCATCCCTGAAGAATC-3’ for shSRGAP2A, 5’-AAGGACAGGCATTGAATATCTTA-3’ for shSRGAP2B/C (targets the 3’UTR of SRGAP2B/C, described in^19^) and shCTNND2#2 inserted at the NheI/BamhI sites. pLV-hsynapsin-CTNND2-myc-WPRE and pLV-hsynapsin-SRGAP2C-HA-myc-WPRE were generated by insertion of the cDNAs at a multiple cloning site (MCS) of a pLV-hsynapsin-MCS-WPRE plasmid (described in^24^) using InFusion cloning (Clontech, Cat#638909).

#### Lentivirus production

HEK293T cells were co-transfected with the lentiviral vector expressing the shRNAs and three plasmids together comprising the lentiviral packaging system. For virus concentration, the viral supernatant was either (1) collected 48h after transfection and ultracentrifuged at 4°C at 25,000 g for 2h; or (2) collected 3 days after transfection and enriched by filter device (Amicon Ultra-15 Centrifuge Filters, Merck, Cat#UFC910008). Viral pellets were resuspended in sterile PBS, aliquoted and stored at –80°C. Titer check was performed for every batch. 100 000 cells were seeded on gelatin-coated coverslips (12-well plates) and infected with 6 serial dilutions of the lentivirus. Medium was changed 24h after and cells were fixed 72 hours after for 1 hour at room temperature with PBS PFA 4%, followed by immunostaining of the relevant marker and Hoechst. From this, the volume of virus needed to infect a given number of cells was estimated (70% for xenotransplantations, 90% of in vitro experiments).

#### shRNA validation

shRNAs were validated in HEK293T cells on exogenously expressed CTNND2 and on endogenous CTNND2 in primary cultures of cortical neurons. Briefly, for validation in a heterologous system, HEK293T cells were co-transfected using Jet-Prime (Polyplus Transfection #POL114-07) according to the manufacturer protocol with either MYC tagged CTNND2 or CTNND2* together with the shRNAs targeting CTNND2 or a control shRNA at a 1:2 ratio. Two days after transfection, cells were collected and lysed in RIPA buffer (150 mM NaCl, 1.0% NP-40, 0.5% sodium deoxycholate, 0.1% SDS, 50 mM Tris, pH 8.0, Sigma-Aldrich) and further processed for western blot analysis of the relative protein expression levels. Conditional knockdown validation was performed in HEK cells by co-expressing pDIO-DSE-mCherry-PSE-MCS containing our shRNAs and pCAG Cre. For the knockdown validation on endogenous *Ctnnd2*, primary cultures of cortical neurons were infected at 4 days in vitro (DIV4) with lentiviruses carrying shCtnnd2#2 or a control shRNA. Infected cultures as well as a non-infected control were harvested at DIV21 and lysed in RIPA buffer under agitation for 1h at 4°C and further processed for western blot analysis (see below).

#### In utero electroporation

In utero electroporation (IUE) was performed as previously described^18,68^. Pregnant Swiss females at E14.5-15.5 (Janvier labs) were anesthetized with isoflurane (3.5% for induction and 1.5-2.5% during the surgery; ISO-VET Piramal Critical Care) and subcutaneously injected with 0.1 mg/kg of buprenorphine for analgesia. The uterine horns were exposed after laparotomy to enable injection of endotoxin-free DNA mix into one ventricle of the mouse embryos. The injected embryos were then electroporated with 4 pulses of 40 V for 50 ms at 500 ms intervals delivered by tweezer-type platinum disc electrodes (5 mm-diameter, Sonidel) and a square wave electroporator (ECM 830, BTX). The volume of injected DNA mix was adjusted depending on the experiments. Plasmids were used at the following concentrations: shRNA vectors: 0.5 µg/µl (adults) or 1 µg/µl (young animals; P10-P23); CTNND2 or shRNA-resistant CTNND2*: 1 µg/µl; SYNGAP-EGFP, SRGAP2C-HA and PSD-95-FingR GFP: 0.7 µg/µl. For conditional knockdown experiments, pCAG-EGFP was co-expressed with pCAG-ERT2CreERT2 and pDIO-DSE-mCherry-PSE-MCS containing our shRNAs (0.5 µg/µl each). For Crispr-Cqs9 knockout experiments, pX330-U6-Chimeric_BB-Cbh-hSpCas9 containing gRNA (0.5 µg/µl) was co-expressed with pCAG-EGFP (1 µg/µl).

#### Slice preparation for confocal imaging

Electroporated mice were sacrificed at the indicated age by terminal transcardial perfusion of 4% paraformaldehyde (Electron Microscopy Sciences) in PBS as described previously^68^. Unless otherwise indicated, 100 μm coronal brain sections were obtained using a vibrating microtome (Leica VT1200S, Leica Micro-systems). Sections were mounted on slides in Vectashield.

#### Primary neuronal cultures

Primary cultures of cortical neurons were performed as previously described^18^ with few modifications. Briefly, mouse cortices from E18.5 embryos were dissected, dissociated and neurons were plated on glass coverslips coated with poly-D-ornytine (80 μg/ml, Sigma) in MEM (Gibco) supplemented with sodium pyruvate, L-glutamine (2 mM) and 10% horse serum. This medium was exchanged 2-3 hours after plating with Neurobasal (Gibco) supplemented with L-glutamine (2 mM), B27 (1X) and penicillin (2.5 units/ml) – streptomycin (2.5 mg/ml). Until use, every 5 days one third of the medium was changed and the cells were maintained at 37°C and 5% CO2. Unless otherwise indicated, all products were from Thermo Fisher Scientific.

#### Immunocytochemistry

After 18 days in vitro (DIV), cultured cortical neurons were fixed for 15 min at room temperature using 4% (w/v) paraformaldehyde in PBS and incubated for 30 min in blocking buffer containing 0.3% Triton X100 and 3% BSA (Sigma) in PBS to permeabilize the cells and block nonspecific staining. Subsequently, cells were incubated for 1h in primary antibody, rinsed extensively, and incubated 45 min in secondary antibodies diluted in blocking buffer. Coverslips were mounted on slides with Vectashield (Vector Laboratories). Primary antibodies were mouse anti-PSD-95 (clone K28/43, Neuromab, 1:200) and rabbit anti-CTNND2 (Abcam #ab184917; 1:500, which was validated in the lab using immunofluorescence on *Ctnnd2*-knocked down cultured mouse cortical neurons). Secondary antibodies were anti-rabbit Alexa-488 (Invitrogen; 1:500) and anti-mouse Cy3 (Jackson ImmunoResearch, 1:500).

#### Confocal image acquisition

Confocal images were acquired in 1024×1024 mode using either Leica TCS SP8 scanning confocal microscope controlled by the LAFAS software and equipped with a tunable white laser and hybrid detectors (Leica Microsystems) or Nikon A1R HD25 scanning confocal microscope controlled by NIS Elements and equipped with an LU-N4/N4S 4-laser unit including 405 nm, 488 nm, 561 nm and 640 nm lasers. On the Leica TCS SP8, we used a 10X PlanApo objective (NA 0.45) to identify electroporated neurons and acquire low magnification images and a 100X HC-PL APO, NA 1.44 CORR CS objective to acquire higher magnification images and z-stacks of dendrites. On the Nikon microscope, we used the following objectives: Plan Apo Lambda 10x Air NA 0.45 (MRD00105) and SR HP Plan Apo Lambda S 100x Silicone NA 1.35 (MRD73950). Z-stacks were acquired with 150 nm spacing and a zoom of x 1.24. Images were blindly acquired and analyzed.

### Ex vivo electrophysiology

#### Slice preparation and electrophysiological recording

Mice were anesthetized with isoflurane (ISO-VET, Piramal Critical Care), decapitated. Brains were quickly removed and placed in ice-cold (4°C) cutting solution containing (in mM): 85 NaCl, 64 sucrose, 25 glucose, 2.5 KCl, 1.2 NaH2PO4, 24 NaHCO3, 0.5 CaCl2, and 7 MgCl2, saturated with 95% O2 and 5% CO2 (pH 7.3–7.4). Acute coronal slices were cut using the 7000 smz-2 vibratome (Campden Instrument). Slices recovered at room temperature in oxygenated artificial cerebrospinal fluid (ACSF) containing (in mM): 125 NaCl, 2.5 KCl, 2 CaCl2, 1 MgCl2, 1.2 NaH2PO4, 24 NaHCO3, and 25 glucose (pH 7.4) for at least 45 min. For electrophysiological recordings, slices were transferred to a submerged recording chamber and continuously perfused at 33–34°C with oxygenated ACSF at a rate of 4 ml/min. Whole-cell voltage and current clamp recordings were acquired from visually identified electroporated or non-electroporated L2/3 CPNs of the somatosensory cortex using an IR-DIC microscope (Olympus) equipped with a 4X objective (UPlanFL, NA 0.13) as well as a 40X water immersion objective (LUMPlan Fl/IR, NA 0.8), 2 automated manipulators (Patchstar, Scientifica) and a ORCAFlash 4.0LT camera (Hamamatsu). Signals were recorded with a Multiclamp 700B Amplifier (Axon instruments), Axon Digidata 1550 (Axon Instruments) and pClamp 10 software (Axon instruments). Borosilicate glass pipettes were pulled to have a resistance of 3-7 MΩ. The recording of mEPSCs and intrinsic excitability was performed using an intracellular solution containing (in mM): 144 K-gluconate, 7 KCl, 10 HEPES, 1 EGTA, 1.5 MgCl2, 2 NaATP, 0.5 NaGTP, (pH adjusted to ±7.3 with KOH). mEPSCs were recorded at a holding potential of –60 mV in the presence of 0.5 µM TTX and gabazine (10 µM). For E/I and AMPA/NMDA ratio recordings, the cutting solution contained (in mM): 83 NaCl, 2.5 KCl, 3.3 MgSO4, 1 NaH2PO4, 22 Glucose, 72 Sucrose, 0.5 CaCl and the extracellular solution contained (in mM): 119 NaCl, 2.5 KCl, 1.3 MgSO4, 1.3 NaH2PO4, 20 Glucose, 26 NaHCO3, 2.5 CaCl supplemented with of 10 μM Gabazine. Cells were recorded with the following intracellular solution (in mM; previously described in Adesnik & Scanziani, 2010): 115 CsMeSO3, 8 NaCl, 10 HEPES, 0.3 NaGTP, 4 MgATP, 0.3 EGTA, 5 QX-314, 10 BAPTA (pH adjusted to ±7.3). A stimulation electrode was placed approximately 100 μm up the apical dendrite and synaptic transmission evoked at 10 s intervals. For the E/I ratio recordings, the stimulation strength was adjusted to evoke a response of approximately 150 pA. The neuron was clamped at –70 mV, the reversal potential of inhibitory synaptic transmission, and subsequently at +10 mV, the reversal potential of excitatory synaptic, to record 10 sweeps of eEPSCs and eIPSCs, respectively. Reversal potentials were validated in a few cells with 10 □M NBQX and 10 □M Gabazine, respectively. The E/I ratio was calculated from the average peak currents or integrated currents (charge) per holding potential. To record intrinsic excitability, cells were held in current clamp mode at resting potential and injected with 1s step currents from –100 pA to +1000 pA with 50 pA increments to assess spike frequency. Current ramps from 0-1000 pA over 1s were used to probe the action potential threshold. In all recordings, access and input resistance were monitored by applying 5 mV hyperpolarizing steps of current. All toxins were obtained from Abcam, salts and other reagents from Sigma Aldrich if not otherwise indicated.

### Xenotransplantation experiments

#### Neonatal xenotransplantation

Neonatal xenotransplantation was performed as already described^47,50^ with some modifications^24^. Human cortical cells that were frozen at DIV25, were thawed and plated on matrigel-coated plates using DDM/B27 and Neurobasal supplemented with B27 (DDM/B27+Nb/B27) medium. Six days after plating (DIV31), cells were dissociated using Accutase and plated on new matrigel-coated plates at high confluency (100,000–600,000 cells/cm^2^) with lentiviral vector. The following day, medium was changed to DDM/B27+Nb/B27 medium. At 14 days after thawing (DIV39), cells were treated with 10μM DAPT (Abcam, Cat#ab120633) for 72 hours. The following day (DIV42), cells were treated with 10μM DAPT and 5μM Cytarabine (ARA-C) (Merck, Cat#C3350000) for 24 hours. The following day (DIV43), medium was changed to DDM/B27+Nb/B27 medium. 19 days after thawing (DIV44), cells were dissociated using NeuroCult dissociation kit (StemCell technologies, Cat#05715) and suspended in the injection solution containing 20 mM EGTA (Merck, Cat#03777) and 0.1% Fast Green (Merck, Cat#210-M) in PBS at 100 000 cells/μl. Approximately 1-2 μl of cell suspension was injected into the lateral ventricles of each hemisphere of neonatal (P0) immunodeficient mice *Rag2*^−/−^ using glass capillaries pulled on a horizontal puller (Sutter P-97).

#### Xenotransplanted mouse cortex processing and immunostaining

Xenotransplanted animals were perfused transcardiacally with ice-cold sucrose 8% PFA 4%. Brains were dissected and soaked in the same fixative overnight, then stored in PBS azide. Then they have been sectioned in 80 μm thickness using vibratome. Slices were transferred into the blocking solution (PBS 0.3% Triton, 5% horse serum, 3% BSA) and incubated for 2 hours. Brain floating slices were incubated 3 days at 4°C in the blocking condition with primary antibody chicken anti-EGFP (1:1,000; ab13970, Abcam). After three PBS washes, slices were incubated overnight at 4°C in PBS with secondary antibody anti-chicken Alexa488 and Hoechst (1:10,000). After three washes in PBS, brain sections were mounted on a slide glass with the mounting reagent (DAKO glycerol mounting medium) using #1.5 coverslips.

#### Image acquisition of human neurons

Confocal images were obtained with Zeiss LSM880 driven by Zen Black and Blue softwares equipped with objectives 10x, 20x, oil immersion 25x and oil immersion 40x & 63x, AiryScan system and argon, helium-neon and 405 nm diode lasers. Specifically, dendritic branches were acquired using an oil immersion x63 objective and AiryScan system with 0.1867357 µm Z steps and 1012×1032 pixel resolution. Neurons located in cortical layers V-VI of the visual and somatosensory cortices have been considered. 1 proximal dendritic branch from each neuron was imaged, located 50 µm from the soma. See below for image acquisition of human xeotransplanted neurons.

### Biochemistry and proteomic experiments

#### Co-Immunoprecipitation in HEK cells

Transfected HEK cells were lysed in 10 mM Tris/Cl pH7.5, 150 mM NaCl, 0.5 mM EDTA and 0.5% Igepal and the protein concentration was determined. From each condition, 0.8-1 mg of total protein was diluted in binding buffer containing 0.05% Igepal, 10 mM Tris/Cl (pH 7.5), 150 mM NaCl, 0.5 mM EDTA and protease inhibitors as well as 25 µl of antibody-conjugated magnetic beads. Depending on the experiment GFP-, MYC-or HA-conjugated magnetic beads were used (GFP-trap and MYC-trap from Chromotek or Pierce anti-HA magnetic beads (Thermo Fisher #88837) respectively). Samples were incubated with the beads for 2 hours at 4°C. After extensive washes, the beads were resuspended in Laemmli buffer and bound proteins were released by boiling (5 min 95°C). Samples were subjected to western blot analysis. Inputs correspond to 15 μg of proteins.

#### Mouse cortex synaptosome protein extraction for western blot analysis

P15 mouse cortices were homogenized in Syn-PER Reagent (Thermo Fisher Scientific, #87793) supplemented with protease and phosphatase inhibitors and centrifuged at 1200 × g for 10 minutes at 4°C. The supernatant was collected and centrifuged 15,000 × g for 20 minutes at 4°C. The resulting pellet was lysed in a buffer containing 100 mM NaCl, 4 mM HEPES pH 7.4, 5 mM EDTA, 5 mM EGTA, 1% CHAPS and inhibitors on a rotating device for 1h at 4°C, then centrifuged at 15,000 x g 10 min at 4°C. The supernatant was collected and protein concentration was measured using BCA assay (Pierce). Samples were boiled for 5 min in Laemmli. Western blotting was performed using precast gels, electrophoresis and transfer chambers from BioRAD according to standard procedures. The following primary antibodies were used: rabbit anti-HA (Cell Signalling #3724S, 1:1,000), rabbit anti-GFP (Thermo Fisher Scientific #A11122, 1:2,000), mouse anti-Myc (Cell Signaling Technology #2276, 1:1,000), mouse anti-CTNND2 (Abcam #ab54578, 1:1,000), rabbit anti-CTNND2 (Abcam #ab184917; 1:1,000), mouse anti-PSD-95 (Neuromab # 75-028, 1:1,000), mouse anti-*α*-TUBULIN (clone DM1A, Merck # 05-829, 1:5,000) and mouse anti-actin (Sigma Aldrich, #MAB1501R, 1:5,000). Donkey anti-mouse or anti-rabbit HRP-conjugated secondary antibodies were used at a 1:10,000 dilution (Jackson Immunoresearch #711-035-150 and #711-035-152 respectively). Protein visualization was performed by chemiluminescence using LumiLight western blotting (Roche) and ImageQuant 800 (GE Healthcare). For western blot on synaptosomes, signals were quantified using Fiji. The signal of each band was normalized to the tubulin signal in the corresponding lane and expressed as a fraction of the average in wild-type samples. All bands shown in in the corresponding figure come from the same membrane.

#### Subcellular fractionation and co-immunoprecipitation for mass spectrometry

Subcellular fractionation was performed from Swiss P15 mouse brains as previously described^68^. All steps were performed at 4°C. Briefly, brains were homogenized in ice-cold HEPES-buffered sucrose (0.32 M sucrose, 4 mM HEPES pH 7.4, 5 mM EDTA, 5 mM EGTA, protease inhibitor cocktail, from Sigma) using a motor driven glass-teflon homogenizer. The homogenate was centrifuged at 3,000 g for 15min. The resulting supernatant was centrifuged at 38,400 g for 15 min, yielding the crude synaptosomal pellet. The pellet was then subjected to hypo-osmotic shock and centrifuged at 38,400 g for 20 min. The resulting pellet was lysed for 1 hour using HEPES-buffered NaCl (100 mM NaCl, 4mM HEPES pH 7.4, 5 mM EDTA, 5 mM EGTA, protease inhibitor cocktail) supplemented with 1% CHAPS (Sigma) and centrifuged at 100,000 x g for 1 hour. The corresponding supernatant is referred to as synaptic fraction or synaptic membranes. Protein concentration was measured using BCA assay (Pierce) and protein samples were subjected to immunoprecipitation. Co-immunoprecipitation were performed using antibodies covalently cross-linked to protein G magnetic beads (Pierce). 36 µg of rabbit anti-CTNND2 or anti-SRGAP2 (directed against the N-terminal^18^, gift from Franck Polleux and Wei Lin Jin) antibodies, or total rabbit IgG in control condition, were incubated for 1h at room temperature and cross-linked with 20 mM DMP (dimethylpimelimidate, Pierce) in 0.2 M Sodium Borate pH 9. After 30 min, the reaction was blocked for 1 hour with 0.2 M Ethanolamine (pH 8). Eventual unbound antibody molecules were washed out by incubating the beads for 5 min in 0.1 M glycine (pH 3). The efficiency of cross-linking was checked by running samples on polyacrylamide 4%–15% gradient gels (Biorad) followed by Comassie Blue staining. 1 mg of total proteins from purified synaptic membranes were diluted in a HEPES-NaCl buffer containing (in mM): 20 HEPES pH 7.4, 150 NaCl, 5 EDTA, 5 EGTA and protease inhibitor cocktail supplemented with 1% CHAPS and incubated overnight at 4°C with antibody-coupled magnetic beads. The beads were rinsed 3 times using HEPES-NaCl buffer supplemented with 0.1% CHAPS and further washed 3 times in a buffer containing 20 mM HEPES (pH 7.4) and 150 mM NaCl. The samples were then subjected to mass spectrometry analysis (see below).

#### Mass spectrometry

Proteomics analysis was performed as previously described^68^. Proteins bound to magnetic beads were washed twice with 100 µL of 25 mM NH4HCO3 and on-beads digested with 200 ng of trypsine/LysC (Promega) for 1 hour in 100 µL of 25 mM NH4HCO3 at room temperature. The peptides were loaded onto an C18 StageTips for desalting and eluted using 40/60 MeCN/H2O + 0.1% formic acid. Mass spectrometry measurement was performed on an Orbitrap Fusion Tribrid mass spectrometer (Thermo Scientific) coupled online to an RSLCnano system (Ultimate 3000, Thermo Scientific) and with a top speed DDA method using higher-energy C-trap collisional dissociation (HCD) fragmentation analyzed in the linear ion trap in rapid mode. The mass spectrometry proteomics data have been deposited to the ProteomeXchange Consortium via the PRIDE^70^ partner repository with the dataset identifiers PXD036488 and PXD036487. For protein identification, data were searched against the Mus musculus (Mouse) UniProt database using Sequest^HT^ through proteome discoverer (version 2.1). Enzyme specificity was set to trypsin and a maximum of two missed cleavage site were allowed. Oxidized methionine, N-terminal acetylation, and carbamidomethyl cysteine were set as variable modifications. Maximum allowed mass deviation was set to 10 ppm for monoisotopic precursor ions and 0.6 Da for MS/MS peaks. The resulting files were further processed using myProMS^71^ v3.9.3. FDR calculation used Percolator and was set to 1% at the peptide level for the whole study.

#### Synaptosome protein extraction from hPSC-derived neurons and western blot analysis

Human cortical cells (frozen at DIV25) were thawed and plated on Matrigel-coated plates using DDM/B27+Nb/B27 medium at 37°C with 5% CO2. Seven days after thawing (DIV32), the cells were dissociated using Accutase and plated on Matrigel-coated plate at high confluency (450,000-700,000 cells/cm^2^). Four days later (DIV36), the medium was changed to DDM/B27+Nb/B27 medium with 10 µM DAPT. Two days after DAPT induction (DIV38), the medium was changed to fresh DDM/B27+Nb/B27 medium. At DIV40, cortical cells were dissociated using NeuroCult Enzymatic Dissociation Kit following manufacturer’s instructions. For L1CAM+ MACS, dissociated cells were incubated with biotin conjugated anti-human CD171(L1CAM) (Miltenyi Biotec, Cat#130-124-046) in MACS buffer at 4°C for 10 min. After washing using MACS buffer, human cells were incubated with anti-biotin microbeads (Miltenyi Biotec, Cat#130-090-485) in MACS buffer at 4°C for 15 min. L1CAM positive selection were carried out with LS columns according to the manufacturer’s instructions. The sorted cells were plated on Poly-L-ornithine-, Laminin– and horse serum-coated 6-well plate at 110,000 cells/cm^2^. The sorted cells were maintained in DDM/B27+Nb/B27 medium. Neurons were infected at DIV45 with lentiviruses, then the medium was changed to DDM/B27+Nb/B27 medium and changed every 3-4 days until DIV70. At DIV70, crude synaptosomes were prepared for each condition from 2 wells with 2M neurons. Cells were homogenized in homogenization buffer (sucrose 1.28 M, Tris(hydroxymethyl)aminomethane 20 mM, MgCl_2_ 4 mM, protease inhibitors) using a plastic cell scraper. Homogenate was spun at 1000 x g for 10 minutes at 4°C. Supernatant was spun at 14,000 x g for 20 minutes at 4°C. Supernatant was kept as the cytosolic fraction. P2 crude synaptosomes were re-suspended in Extraction Buffer (HEPES 50mM pH = 7.5, NaCl 150mM, EDTA 2mM, NP-40 1%, Triton 0.5%, Na_3_VO_4_ 1mM, NaF 30mM, protease inhibitors) and extracted for 1 hour, then centrifuged at 10,000 x g for 30 minutes at 4°C to pellet insoluble material. Subsequently, samples were diluted to final 1x Laemmli buffer at 95°C for 5 min. Protein extracts from human synaptosomes were separated using NUPAGE 4-12% Bis-Tris Protein Gel at the voltage of 90 V for 2 hours in MOPS buffer and then transferred to PVDF blotting membrane at the voltage of 100 V for 100 minutes. The membrane was blocked in the buffer (5% skim milk and 0.1% Tween20 in TBS) for 1 hour at room temperature and subsequently incubated in the blocking buffer containing rabbit anti-SYNGAP1 (1:5000; ThermoFisher #PA1-046), rabbit anti-SRGAP2A (1:5000; Abcam # ab124958), and mouse anti-synaptophysin (1:5000; Synaptic Systems #101011) overnight at 4°C. The membrane was then incubated with secondary antibody anti-Rabbit and Mouse IgG antibody conjugated with HRP in blocking solution at room temperature for 1 hour. SuperSignal™ Western Blot Substrate were used for signal detection. The CTNND2 staining (rabbit anti-CTNND2; 1:5000; Abcam ab184917) was performed similarly but after blot stripping for 15 minutes using Restore™ PLUS Western Blot Stripping Buffer (ThermoFisher #46430). Signals were measured using Fiji, normalized synaptophysin and expressed as ratio of mean shScramble value. All bands shown in the figure are from the same membrane. Note that these data were generated from the same series of experiments as in^24^.

#### Subcellular fractionation and co-immunoprecipitation using human brain samples

Crude synaptosome extracts were prepared from thaw biopsy homogenized using a Dounce homogenizer in Homogenization buffer (sucrose 1.28 M, Tris(hydroxymethyl)aminomethane 20 mM, MgCl_2_ 4 mM, protease inhibitors). Homogenate was spun at 1000xg for 10 minutes at 4°C. Supernatant was spun at 14,000 x g for 20 minutes at 4°C. Supernatant was kept as the cytosolic fraction. P2 crude synaptosomes were re-suspended in Extraction Buffer (HEPES 50mM pH = 7.5, NaCl 150mM, EDTA 2mM, NP-40 1%, Triton 0.5%, Na_3_VO_4_ 1mM, NaF 30mM, protease inhibitors) and extracted for 2 hours and centrifuged at 10000xg for 30 minutes at 4°C to pellet insoluble material. Samples were incubated overnight on a wheel in a cold room with protein A magnetic beads coupled with 1ug of rabbit anti-SRGAP2A or 1ug of rabbit anti-CTNND2 or 1ug of rabbit IgG (ThermoFisher) antibodies. Beads were washed 4 times with the washing solution (HEPES 50 mM pH = 7.5, NaCl 150 mM, EDTA 2 mM, NP-40 1%, Triton 0.5%, Na_3_VO_4_ mM, NaF 30 mM) and once with PBS. Subsequently, samples were eluted in 2x Laemmli buffer at 95°C. The input (in 1x Laemmli buffer) and immunoprecipitated samples were run in NUPAGE 4-12% Bis-Tris Protein Gel at the voltage of 90V for 2 hours in MOPS buffer and then transferred to PVDF Blotting Membrane at the voltage of 100V for 100 minutes. The membrane was blocked in the buffer (5% skim milk and 0.1% Tween20 in TBS) for 1 hour at room temperature and subsequently incubated in the blocking buffer containing rabbit anti-SYNGAP1, rabbit anti-CTNND2 or rabbit anti-SRGAP2A antibodies overnight at 4°C, followed by the incubation in the blocking solution containing secondary antibody anti-Rabbit IgG antibody conjugated with HRP at room temperature for 1 hour. Pierce ECL Western Blotting Substrate was used for signal detection. Note that these data were generated from the same series of experiments as in^24^.

## QUANTIFICATION AND STATISTICAL ANALYSIS

### Confocal image analysis

*Mouse samples:* In immunocytochemistry experiments on cultured neurons, the association of CTNND2 with PSD-95 was analyzed using ICY v1.9.9.1^72^. Regions of interest were manually drawn around dendrites and clusters in each channel were detected using the spot detector plugin^73^ of ICY. Then the association between channels was assessed with SODA 2-channel colocalization protocol^74^ using the synaptic markers as reference channel (radius set to 1.5x the max. feret diameter). The fraction of PSD-95 clusters associated with a cluster of CTNND2 was defined as association index. In brain slices, dendritic spine density was quantified in oblique dendrites originating from the apical trunk using Fiji^75^ (https://fiji.sc/). Only dendrites that were largely parallel to the plane of the slice and acquired from sections of comparable rostro-caudal position were analyzed (no more than 1 dendrite per neuron). The density of dendritic spines along dendrites was calculated as described before^18,19^ over a minimal dendritic length of 50 μm starting from the object closest to the branchpoint with the main apical dendrite (minimum 100 spines per dendrites). The length of the dendritic segment was measured on the z projection. For conditional knockdown experiments, only cells expressing EGFP and mCherry, whose expression was induced by recombination after tamoxifen exposure were analyzed and spines were quantified based on EGFP fluorescence. To assess SYNGAP1 and PSD-95 levels in spines, ROIs were manually drawn around dendrites and clusters were detected using the ICY spot detector plugin. Fluorescence intensity associated with SYNGAP1 and PSD-95 in each spine was normalized to mean local filler intensity and compared between conditions. Since CTNND2-GFP signals were located at the periphery of the spines and CTNND2 was also diffuse in the shaft, fluorescence intensity associated with CTNND2 in spines was normalized to CTNND2 fluorescence in the dendritic shaft. One dendrite per cell was analyzed. All images were blindly analyzed.

*Xenotransplantation experiments:* morphometric analyses of dendritic arbour and dendritic spines were performed by manual 3D reconstruction using Imaris software (Bitplane). Dendritic spines were quantified in proximal dendrites located 50 µm from the soma. Spine density was defined as the number of quantified spines divided by the length over which the spines were quantified.

### Electrophysiological data analysis

We compared electrophysiological properties in electroporated CTNND2-depleted neurons versus neighboring non-electroporated control neurons. Voltage and Current clamp data were sampled at 10 kHz and filtered at 2 kHz. All traces were analyzed using pClamp 10.0 (Molecular Devices). Miniature currents were analyzed over 1 min-periods. Overlapping events were excluded from amplitude analysis. The AMPA/NMDA was defined as the peak of the EPSC at – 70 mV over the amplitude of the EPSC at +40 mV 50 ms after stimulation (5 sweeps per holding potential). Cells showing >20% change in access and input resistance upon application of 5 mV hyperpolarizing steps of current were excluded from the analysis.

### Proteomics analysis: label-free quantification

Label free quantification was performed by peptide Extracted Ion Chromatograms (XICs) computed with MassChroQ version 2.2.1^76^. For protein quantification, XICs from proteotypic and non-proteotypic peptides shared between compared conditions (TopN matching), and all charge states and sources were used. Global MAD or median and scale normalization was applied on the total signal to correct the XICs for each biological replicate (n = 3). To estimate the significance of the change in protein abundance, a statistical test based on a linear model adjusted on peptides and biological replicates was performed and *p*-values were adjusted with a Benjamini–Hochberg

FDR procedure. Gene ontology analysis was performed using the SYNGO database (https://syngoportal.org/)^35^ using brain expressed genes as background. The genes annotated in SYNGO were further analyzed using g:Profiler (https://biit.cs.ut.ee/gprofiler/gost) for human phenotypes^77^.

### Statistical analysis

Statistical analyses were performed with Prism 7 (GraphPad Software). Data are obtained from a minimum of three independent experiments. For in utero electroporations data were obtained from at least three experiments or three animals from two independent litters. For statistical analysis, normality of the distributions was assessed using the D’Agostino-Pearson normality test. In case of normal distributions, we used two-tailed Student’s t test or one-way analysis of variance followed by Tukey’s post hoc test. Non-normal distributions were assessed using the non-parametric Mann-Whitney test. In boxplots, whiskers correspond to the minimal and maximal values. Other data represent mean ± SEM. Sample size and statistical tests are reported in each main and supplementary figure. The significance threshold was placed at 0.05 (NS = p > 0.05; *= p < 0.05, **= p <0.01; ***= p<0.001).

## SUPPLEMENTAL INFORMATION

Document S1. Figures S1–S6

Table S1

