## Supplementary information for "CTNND2 regulation by the SRGAP2 protein family links human evolution to synaptic neoteny"

### SUPPLEMENTAL INFORMATION

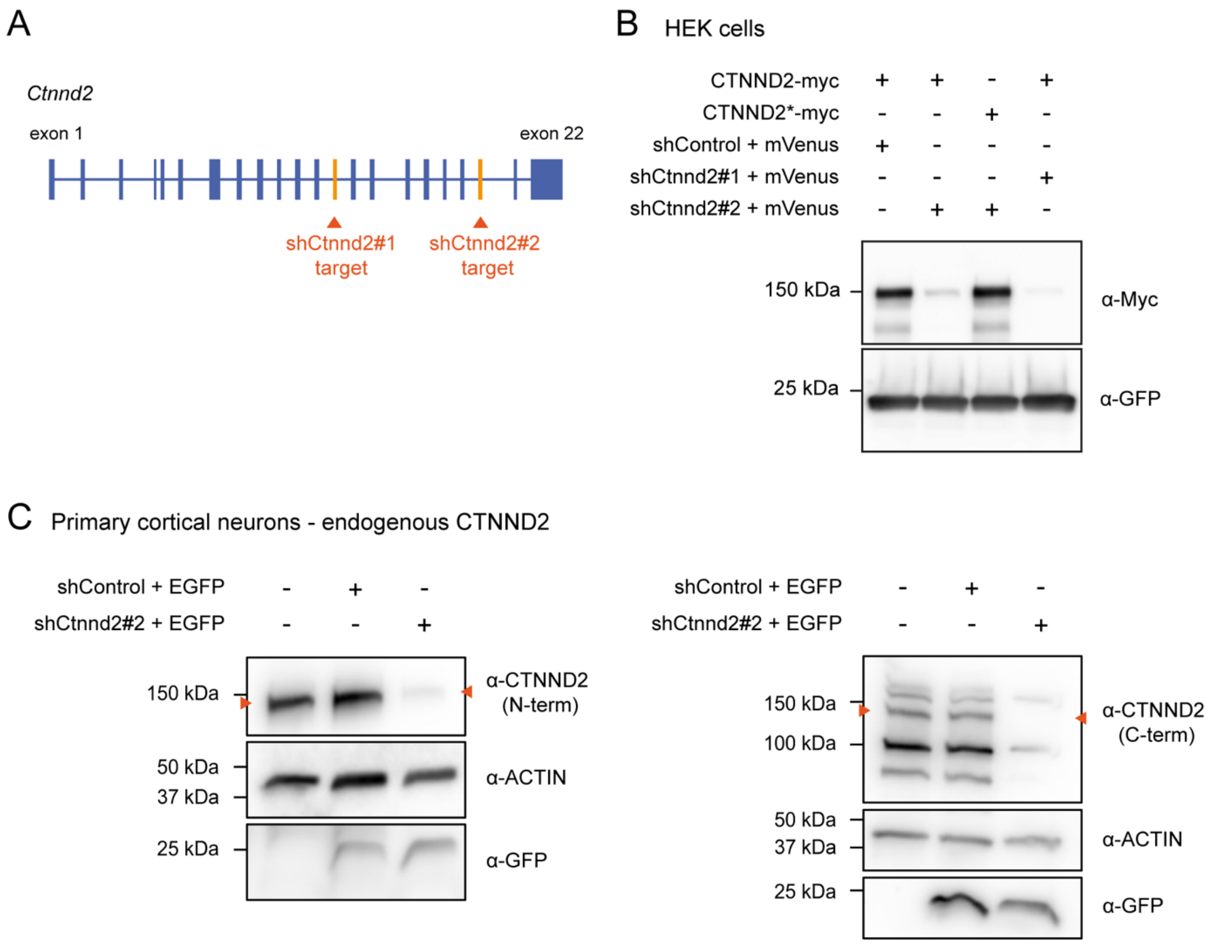


**Figure S1 Validation of shRNAs against *ctnnd2* and rescue construct.** Related to Figures 2-7

(A) Gene model and main cortical isoform of *ctnnd2* (adapted from Gtex portal data obtained on 13/07/2022). Exons containing the sequences targeted by the shRNAs are highlighted in orange.

(B) Validation of shRNAs in HEK cells and primary cortical neurons using western blot. HEK cells were transfected with indicated cDNAs and harvested after 48 hours. CTNND2* contains silent point mutations that make it resistant to shCtnnd2#2 (rescue construct). shControl: scrambled control shRNA. $\alpha$ represents “antibody against”.

(C) Cortical neurons were infected at DIV 4 with lentiviruses driving the expression of shRNAs along with EGFP and then collected at DIV 21. CTNND2 was detected using an antibody directed against the N-terminal (left) or C-terminal (right) region of the protein respectively.

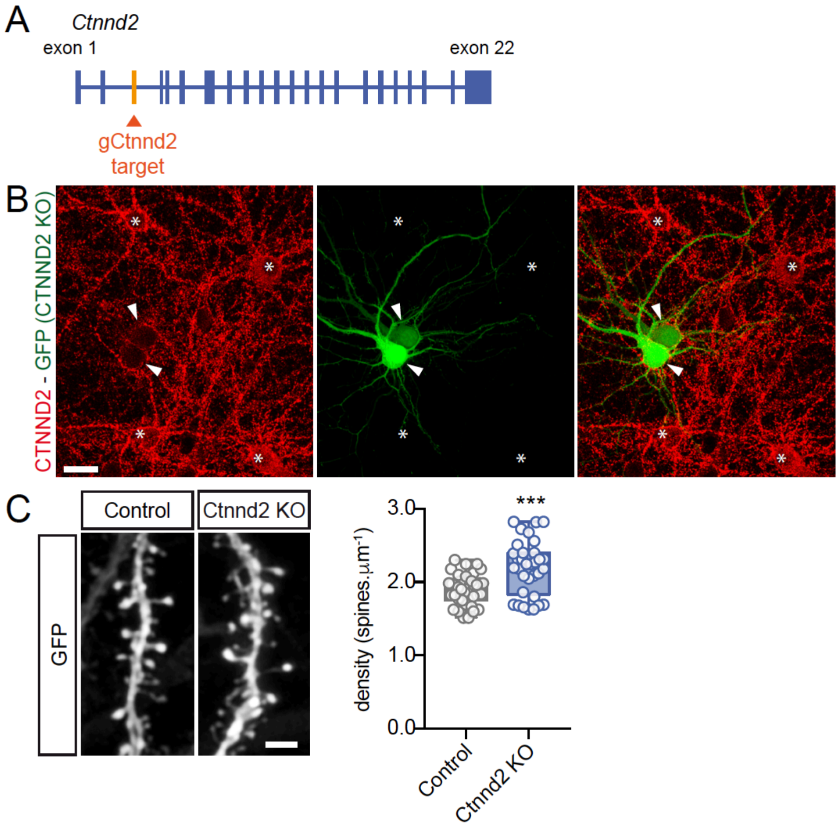


**Figure S2 CRISPR-Cas9-mediated knockout of *ctnnd2* increases spine density**.
Related to Figure 2
(A) Exon-intron structure of *ctnnd2*. The gRNA used in knockout experiments targets exon 3 (gCtnnd2).
(B) Validation of CTNND2 KO. L2/3 CPNs were electroporated ex utero at E15.5 with pCAG GFP along with a plasmid driving the expression of CAS9 and gCtnnd2. Cortical neurons from the electroporated hemisphere were then dissociated, plated on glass coverslips and cultured for 17 days in vitro before fixation and immunostaining of endogenous CTNND2. Note that CTNND2 is detected in wild-type neurons (asterisks) but not in CTNND2 KO neurons (arrowheads). Scale bar: 20 µm.
(C) Representative segments of P21 dendrites and quantification of spine density in neurons expressing spCas9 and gCtnnd2 (Ctnnd2 KO) or the spCas9 alone (Control) along with soluble EGFP to visualize dendritic spines. Scale bar is 2 µm. $N_{Control}$ = 39 (5); $N_{gCtnnd2}$ = 33 (6). Individual datapoints represent single dendrites (one dendrite per cell was quantified). Mann-Whitney test, ***: $p < 0.001$.

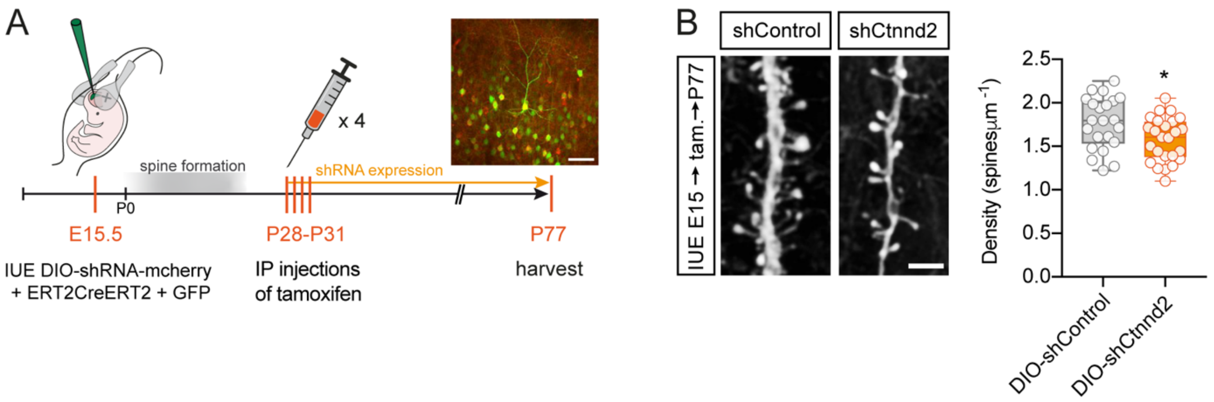


**Figure S3 Induced *Ctnnd2* knockdown from P28 decreases spine density in adults**.

Related to Figure 2

(A) Experimental workflow of *Ctnnd2* conditional knockdown experiments. All electroporated neurons are GFP-positive. ShControl or shCtnnd2 and mCherry expression is induced after tamoxifen injections between P28 and P31. Scale bar 50 μm.

(B) Representative segments of dendrites and quantification of spine density in conditional knockdown experiments. $N_{(control)}$ = 22 (3), $N_{(shCtnnd2)}$ = 25 (4). Statistics: *: $p < 0.05$, Mann-Whitney test.

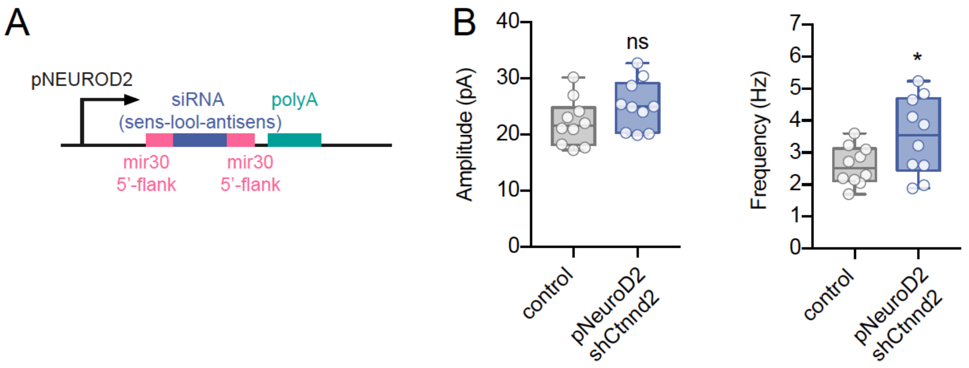


**Figure S4 *Ctnnd2* knockdown in postmitotic neurons increases mEPSCs frequency.**
Related to Figure 3
(A) Diagram of RNAi vector (gift from Juliette Godin, IGBMC, France) to knockdown *Ctnnd2* in postmitotic neurons.
(B) Quantification of mEPSC recordings in L2/3 CPNs of young (P10-P12) mice following IUE with a vector expressing shCTNND2 under NEUROD2 promoter (described in A). Data points in boxplots represent mean values for individual cells. N = 10 cells (from 3 mice) in each codition. Statistics: *: $p < 0.05$, ns: $p > 0.05$; Student t-test.

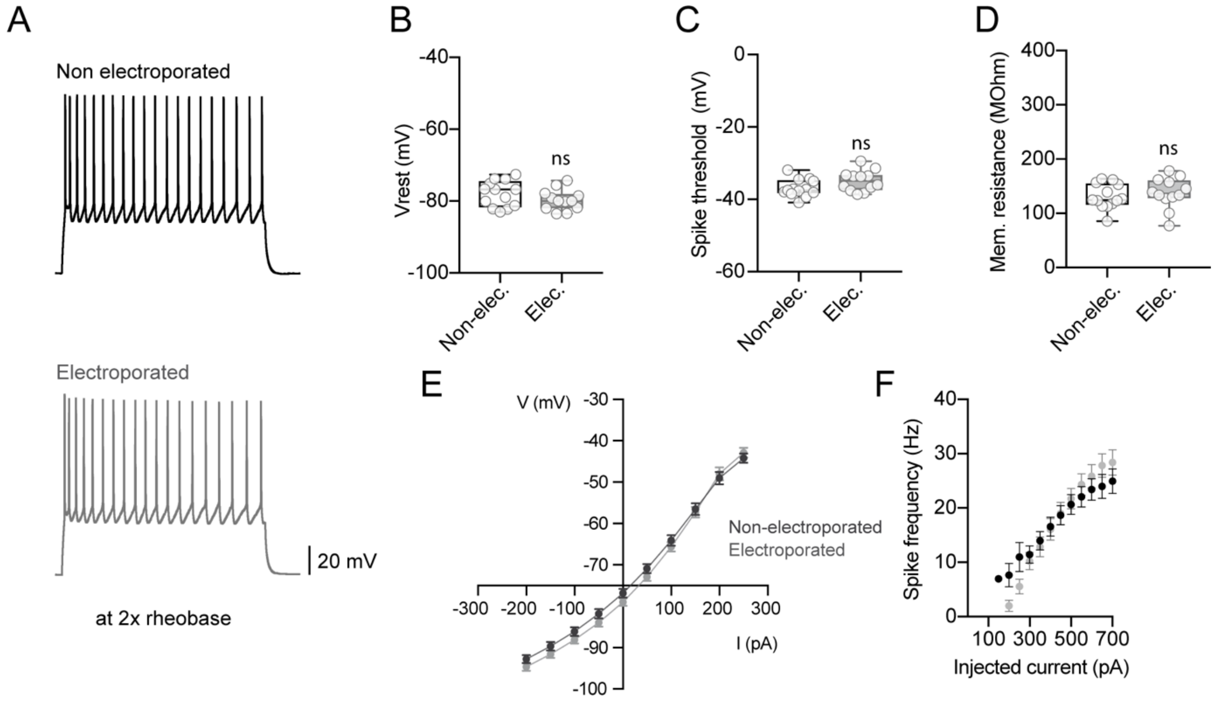


**Figure S5 In utero electroporation does not affect intrinsic neuronal properties.**

Related to Figure 4

(A) Traces of action potentials evoked at twice the rheobase by 1s depolarizing current steps in whole-cell current-clamp recordings.

(B-F) Quantification of intrinsic membrane properties in non-electroporated neurons (non-elec.) and neurons electroporated (elec.) with a plasmid expressing shControl and tdTomato. $N_{control}$= 13 cells (5 mice), $N_{electroporated}$ = 12 (5). ns: $p > 0.05$, Student t-test.

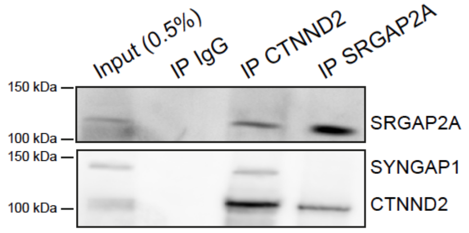


**Figure S6 SRGAP2A, CTNND2, SYNGAP1 synaptic complex.**
Related to Figure 7.
Synaptosomal protein extract from a human temporal cortex biopsy (45-year-old) with control (IgG), CTNND2 and SRGAP2A immunoprecipitation (IP) followed by staining for CTNND2, SRGAP2A and SYNGAP1 (SRGAP2A IP: 3 ; CTNND2: n=2 independent experiments). Note that SRGAP2A and CTNND2 co-immunoprecipitate. SYNGAP1 co-immunoprecipitates with CTNND2 but not SRGAP2A.
